# Diversity, structure-function relationships and evolution of cell wall-binding domains of staphylococcal phage endolysins

**DOI:** 10.1101/2025.02.06.636829

**Authors:** Roberto Vázquez, Diana Gutiérrez, Bjorn Criel, Zoë Dezutter, Yves Briers

## Abstract

Endolysins, encoded by phages, lyse bacterial hosts at the end of the replication cycle by degrading peptidoglycan. Consequently, they have evolved in response to host cell wall structures, leading to complex modular architectures, particularly in Gram-positive bacteria. These architectures feature diverse enzymatically active domains (EADs) and cell wall-binding domains (CBDs).

This study investigates the structure-function relationships of CBDs in staphylococcal phage endolysins, exploring their evolutionary origins and the extent to which binding specificity can be predicted from sequence data. A set of 182 staphylococcal endolysin sequences was analyzed, revealing predominantly three-domain architectures, occasionally disrupted by species-specific mobile genetic elements. Most CBDs exhibited an SH3-like fold, classified into two major subfamilies: SH3b_P (including the well-characterized SH3_5 family) and SH3b_T. The composition of endolysin domains correlated with specific CBD families, suggesting co-evolution of CBDs and compatible EADs to ensure functional synergy.

To assess binding properties, 24 CBDs were fused to eGFP and tested against a panel of staphylococci, revealing diverse specificity profiles. However, no clear correlation emerged between binding specificity, phylogenetic subgroups, or bacterial hosts. This suggests that minor structural modifications significantly impact function and that CBD specificity is not a major selective pressure in the staphylococcal bacteria-phage interface.

## 1. INTRODUCTION

In the last years, the antibiotic resistance crisis has sparked interest in a number of non-conventional therapeutic agents to treat bacterial infections [1,2]. One of such alternative antibacterials are phage endolysins. These enzymes are encoded by (bacterio)phages and disrupt the peptidoglycan layer in the bacterial cell wall at the end of the phage replication cycle. Particularly, all known dsDNA phages bear at least an endolysin that acts upon the maturation of the new virions inside the bacterial host cell, enabling the release of those viral particles and lysing the host in the process. A remarkable body of work has been generated in the last couple of decades, demonstrating the antibacterial capacity of exogenously applied endolysins against pathogenic bacteria [3–6]. Endolysins acting against Gram-positive bacteria are typically modular, *i.e.*, they are composed of different domains devoted either to the catalytic activity itself (enzymatically active domains, EADs) or to bind the cell wall of the host (cell wall-binding domains, CBDs), and these different domains are found in different architectural combinations among the natural endolysin diversity [7,8]. Since long, it is accepted that phage genomes dynamically evolve in a modular manner. Indeed, phage genomes are mosaic, composed by specific modules that are exchanged within the bacteria-phage interplay [9]. The domains of endolysins also take part in this natural recombination process [10], and thus, inspired by nature, researchers have been using them as modules in combinatorial engineering to render novel domain combinations that are better adjusted to given applied objectives [11,12]. While this has substantially enriched the application potential of endolysins, it is also true that combinatorial engineering remains largely empirical, therefore heavily relying on *ad hoc* high-throughput technical solutions [13]. Even with these advances it is practically impossible to analyze the full modular design space. Thus, a deeper insight into the structure-function relationships of phage lysin modules would provide, on the one hand, a better understanding on the evolutionary constraints and mechanisms that have (transiently) fixed certain architectural variants in the current lysin diversity, and, on the other hand, a knowledge base to conduct a more rationalized and efficient lysin engineering. With this in mind, the general aim of this work is to characterize the structural diversity of a set of endolysin modules and their linkage to functional behavior. We chose the endolysins from staphylococcal phages, focusing on their CBDs, as these domains are expected to have the greatest impact on specificity [14].

The genus *Staphylococcus* was selected because it has traditionally been the primary subject of lysin engineering, given its importance as a human and veterinary pathogen and, moreover, the concerning surveillance data available on its drug resistance and global mortality [15]. For example, *Staphylococcus aureus* is the pathogen for which the first phase 2 and 3 trials have been performed [16]. Thus, there is already a large research corpus on which to build new knowledge. Particularly, it is well-known that anti-staphylococcal endolysins, as most of the endolysins found in phages that infect Gram-positive hosts, typically have an architecture with two putative EADs at the N-terminal end of the protein plus a CBD at the C-terminal end [7,17]. This CBD is usually predicted to belong to the *SH3_5* family (PF08460) although in many other cases no known family has been assigned yet, or their family has only been recently described, as is the case of *SH3b_T* (PF24246) [14]. The *SH3_5* family itself belongs to the wider group of SH3b domains, which, in turn, are the type of SH3 folds that are found in bacterial proteins (and the viruses that infect them). SH3 folds are ubiquitous in nature, and they are one of the oldest and simplest protein folds, consisting of a barrel-like β-sheet structure, which is known to work as a binding domain in general [18]. SH3 domains have been shown to be involved in protein:protein contacts, subcellular localization and, of course, binding epitopes at certain sites, such as the bacterial cell wall [19]. Specifically, some of the SH3b domains found in endolysins from different hosts bind either directly certain peptidoglycan chemotypes (as is the case for the experimentally tested *SH3_5* domains in anti-staphylococcal lysins) or other surface ligands such as the teichoic acids [20,21].

To address the proposed question of this study, we devised a workflow comprising: (i) the compilation of a sequence set of anti-staphylococcal endolysins; (ii) mining the known and (potentially) unknown CBDs from such sequences; (iii) analyzing their structural nature in connection with the architectures in which they appear; and (iv) generating a subset of reporter:CBD fusion proteins to be tested against a collection of staphylococcal strains. As such, we aimed to gain a structured understanding of the diversity of CBDs and to link it to empirical evidence on their binding specificity at a systematic level.

## 2. MATERIALS AND METHODS

### 2.1. Mining staphylococcal CBDs

A total of 816 endolysin sequences with a staphylococcal or non-annotated host were initially retrieved from PhaLP (version PhaLP_v2020_06). The sequences were pairwise aligned using the ClustalOmega algorithm and clustered based on a similarity distance matrix (**Supplementary Table S1**). Only those sequences with non-annotated hosts that lied within the visually distinct clusters in which there were representatives annotated as staphylococcal phages were subsequently kept (which corresponded to an inclusion criterium of >37.32% similarity to any already included sequence). The sequence fragments predicted to be CBDs (*i.e.*, belonging to a Pfam family that are known CBDs, such as *SH3_5* or *PG_binding_1*) according to metadata from PhaLP plus unannotated stretches with a length >40 amino acids were extracted as putative CBDs. Such putative CBDs were manually curated to remove those sequence fragments belonging to incomplete endolysins (due *e.g.* to a genomic insertion, see later **Figure 3** and its discussion) or misannotated structural lysins. After this curation, a final set of 182 sequences was obtained (**Supplementary Table S2**). An initial guiding UPGMA phylogenetic tree was built with the CBD sequences obtained from domain predictions or unannotated sequence regions. According to such tree, 40 entries representative of the full variability were selected to perform three-dimensional structure predictions. Based on such predictions, the delineation of the CBDs was improved to proceed with further analyses, including all amino acids participating in secondary structures within the domain. As a part of this delineation, the CBDs that consisted of more than one domain repeat were also separated in their individual structural units to make them more comparable to their single-repeat counterparts in the analyses.

### 2.2. Bioinformatic analyses

The amino acid sequences of the delineated CBDs were aligned using the ClustalOmega algorithm as implemented in R package “msa” [22]. Sequence identity-based distance matrices were obtained from the alignments using “seqinr” library [23]. Then distance-based trees were generated using the UPGMA algorithm implemented in the library “phangorn” [24], and visualized using “ggtree” [25]. The tree was further annotated with different categories: (i) structural groups were obtained from the (re)classification of CBD families as explained in **Figure 2**; (ii) host species were retrieved together with the mined endolysin sequences; (iii) phylogenetic clusters were obtained by hierarchical clustering followed by manual curation to remove clusters with less than 7 elements (but explicitly withholding the 2-element cluster containing the only examples of CBDs belonging to *PG_binding_1* family); (iv) the architectures were annotated based on Pfam domain predictions on the full lysin sequences. For the latter annotation, the genomic context of each lysin was also considered, since it is known that many staphylococcal endolysin genes are interrupted by self-splicing introns. For those endolysin entries for which an intron and further domains pertaining to the final endolysin product were detected in the genome, the architecture was described considering said final, full product, and the elements of the endolysin that are not present in the CDSs acquired from PhaLP are included between square brackets in **Figure 1** and **Supplementary Table S2**. For example, UniProt entry H9A141 (**Supplementary Figure S2**) contains only the remnants of an Amidase domain plus an *SH3_5* CBD, but upstream of the gene encoding for H9A141, a second CDS coding for a single *CHAP* domain can be found, followed by a plausible self-splicing intron (which contains an *HNH* endonuclease, see below for discussion). Therefore, the architecture of H9A141 is described as [CHAP]:[intron]:Amidase:SH3b.

**Figure 1.**
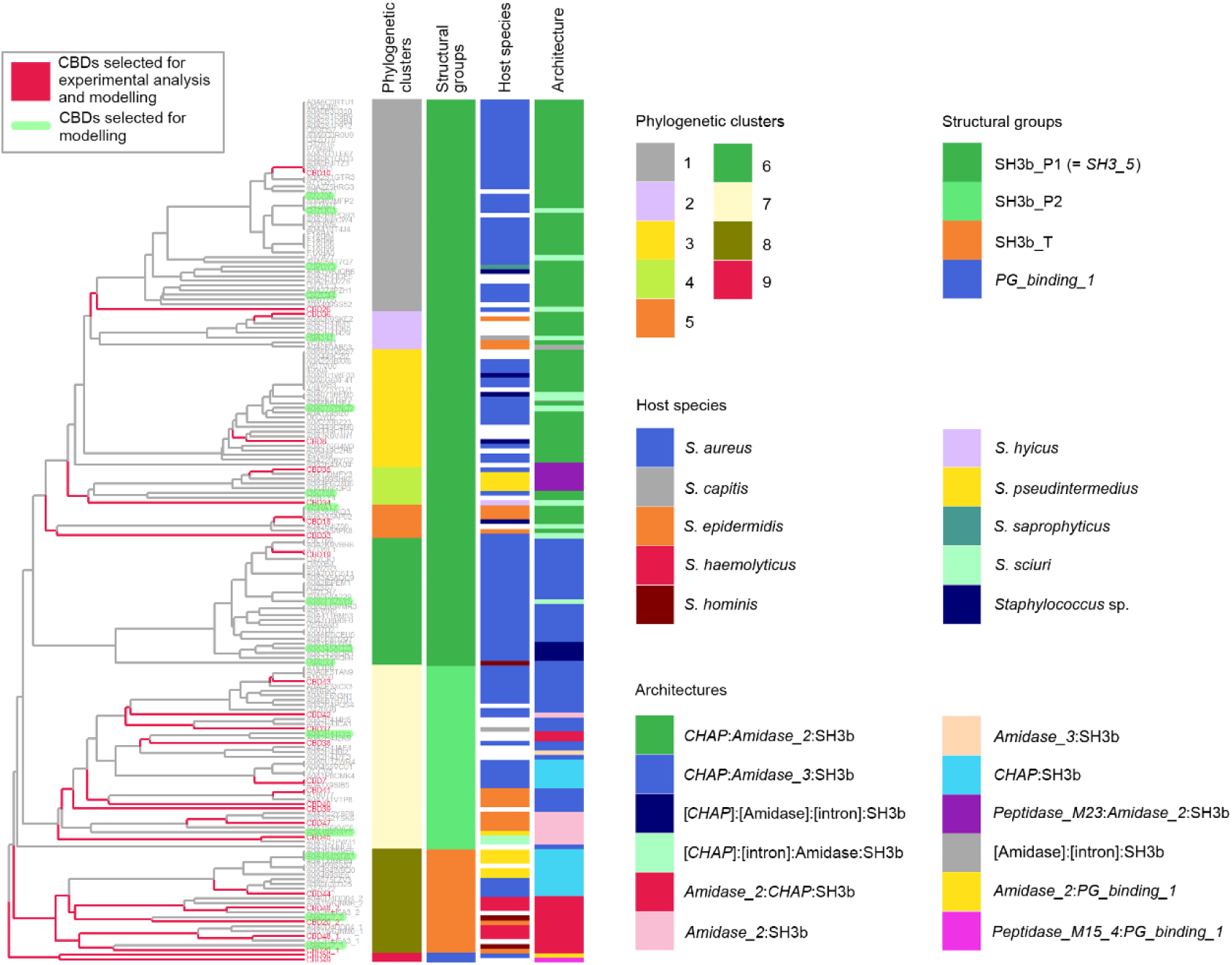
Phylogenetic tree of the proposed individual CBD repeats found among the staphylococcal lysins extracted from the PhaLP database. The tips of the tree are labelled either with the UniProt accession number of the original endolysin (plus ‘_1’ or ‘_2’ in those endolysins containing two CBD repeats) or with the ID of the corresponding tile (CBD10, CBD26, etc., see **Table 2**). Metadata on the custom phylogenetic clusters, inferred structural groups, host species annotated in the source phage genome and the predicted architecture of each endolysin are depicted in the bars adjacent to the tree according to the color legend. Architectural elements found in adjacent genes at the source genome are shown between square brackets.

**Figure 2.**
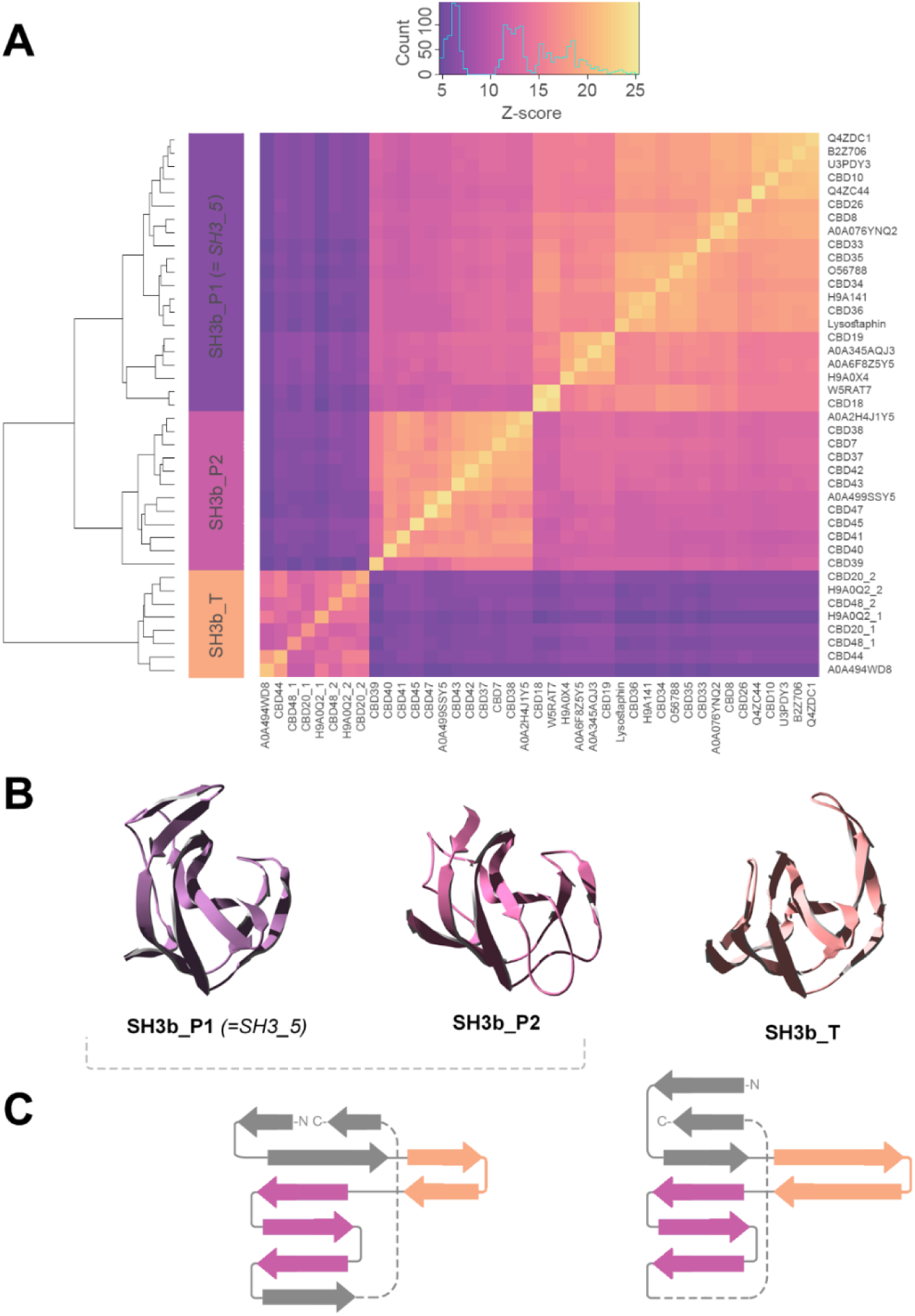
Structural classification of SH3b CBDs from anti-staphylococcal endolysins. (A) Pairwise similarity matrix based on a DALI 3D structure comparison of the structural predictions obtained for a selected subset of 42 representative CBD repeats (Figure 1) plus lysostaphin CBD. (B) Representative three-dimensional structure models of the three large SH3b structural groups uncovered by the structural similarity analysis (from left to right: CBD8, CBD42 and the second repeat of CBD20). (C) Proposed differential topologies for *SH3b_P* (left) and *SH3b_T* (right) as observed in the analyzed set of 3D models.

### 2.3. Three-dimensional structure prediction

The three-dimensional structure of the 42 selected CBDs were predicted using AlphaFold2 with the MMseqs2 search engine as implemented in ColabFold [26], with default parameters. The pLDDT values for the best predictions per protein are available in **Supplementary Table S2** (mean pLDDT was 87.9, ranging between 79.3 and 96.6). The structural models were explored and rendered using Swiss-PDBViewer [27]. DALI server was used to structurally align the trimmed CBDs (available as **Supplementary File S1**) and obtain a pairwise structural similarity matrix and structural conservation plots [28].

### 2.4. Bacterial strains and culture conditions

*Escherichia coli* TOP10 was used for cloning and *E. coli* BL21(DE3) for protein expression, both cultured in lysogeny broth (LB) at 37 °C with shaking (200 rpm) or in LB plates with 2 % (w/v) bacteriological agar. Ampicillin at a final concentration of 100 μg/ml was used as a positive selection marker for plasmid pVTE while 50 μg/ml kanamycin was added to positively select for pVTD3 *E. coli* transformants. 5% (w/v) sucrose was used for any of the two plasmids to negatively select against vectors lacking insertion of the heterologous gene constructions, as previously described for the VersaTile method [13]. Staphylococci (**Table 1**) were cultured in tryptic soy broth (TSB) at 37 °C with shaking (200 rpm) or on TSB plates containing 2 % (w/v) bacteriological agar. Bacterial stocks were made by adding 20% (v/v) glycerol to grown bacterial cultures and were kept at −80°C.

**Table 1.** Staphylococcal strains used in this work.

| Species | Strain | Source |
| --- | --- | --- |
| *S. aureus* | Sa9 | [29] |
|  | ATCC 15981 | [30] |
|  | MRSA-17 | Donated by Chris Callewaert, skin isolate |
|  | 119C | Donated by Chris Callewaert, skin isolate |
| *S. epidermidis* | F12 | [31] |
|  | Z2LDC14 | [31] |
|  | 25 | Donated by Chris Callewaert, skin isolate |
|  | CC28 | Donated by Chris Callewaert, skin isolate |
| *S. hominis* | CC40 | Donated by Chris Callewaert, skin isolate |
|  | 88F | Donated by Chris Callewaert, skin isolate |
|  | 119C | Donated by Chris Callewaert, skin isolate |
|  | 96 | Donated by Chris Callewaert, skin isolate |
| *S. capitis* | SK14 | Donated by Chris Callewaert, skin isolate |
|  | VCU116 | Donated by Chris Callewaert, skin isolate |
|  | CC44 | Donated by Chris Callewaert, skin isolate |
| *S. haemolyticus* | Shae 2 | Donated by Chris Callewaert, skin isolate |
|  | ZL89-3 | [32] |
| *S. sciuri* | DB156 | Donated by Chris Callewaert, skin isolate |
|  | DB258 | Donated by Chris Callewaert, skin isolate |
| *S. hyicus* | DB222 | Donated by Chris Callewaert, skin isolate |
|  | DB171 | Donated by Chris Callewaert, skin isolate |
| *S. saprophyticus* | 36 | Donated by Chris Callewaert, skin isolate |
| *S. xylosus* | ZL61-2 | [32] |
| *S. kloosi* | ZL74-2 | [32] |

### 2.5. Plasmid construction and DNA manipulation

Unless already present in the UGent VersaTile repository, synthetic gene fragments encoding the CBDs to be tested in this work plus the flanking regions needed for VersaTile cloning [13] were purchased from Twist Bioscience (South San Francisco, CA, USA). Such genes were inserted in VersaTile entry vector pVTEIII by digestion with *SapI* plus ligation. The ligation products were subsequently used for transformation of *E. coli* TOP10 by electroporation and recombinant transformants were selected by plating on LB plus ampicillin and sucrose. The TOP10 cells were used as a source for “tiles” (gene fragments flanked by position markers and BsaI recognition sites for subsequent Versa-Tile assembly), which were all confirmed by Sanger sequencing (LGC Genomics, Teddington, UK) and stored at -80°C.

The eGFP-CBD fusion coding sequences were obtained by assembling an eGFP coding sequence at the first position, the corresponding CBD at the second position and a 6×histidine tag plus stop codon at the third position. The used position markers were NCCATG and NCAGGN, NCAGGN and NGAAGN, and NGAAGN and NAGTAN, respectively for positions 1, 2, and 3. Restriction/ligation reactions for insertion into the destination vector were conducted using BsaI as the type IIs restriction enzyme, as described before (13). *E. coli* BL21(DE3) was transformed with the final constructs, selecting the transformants with non-empty pVTD3 vector with kanamycin and sucrose, and their sequence was verified by Sanger sequencing. A list of the CBD tiles used in this work, including their source NCBI entry and the delineation coordinates can be found in **Table 2**.

**Table 2.** CBD tiles used in this work.

| Tile ID | Tile name | Source (Uni-Prot) | Delineation (aa start:end) | Phage annotated host |
| --- | --- | --- | --- | --- |
| CBD7 | Romulus175_SH3 | M4I0E5 | 186:252 | *S. aureus* |
| CBD8 | LysRODI_SH35 | A0A0D3MUT0 | 406:496 | *Staphylococcus* sp. |
| CBD10 | LysH5_SH35 | B7T0F1 | 397:486 | *S. aureus* |
| CBD18 | LysC1C_SH35 | A0A0D3MVQ2 | 381:484 | *Staphylococcus* sp. |
| CBD19 | LysA72_SH35 | B7T0L1 | 395:484 | *S. aureus* |
| CBD20 | LysIPLA5_SH3 | I6T7G5 | 380:574 | *S. epidermidis* |
| CBD26 | PlyGRCS SH35 | W6E9K2 | 151:250 | *S. aureus* |
| CBD33 | PlyBS2_SH35 | A0A2R3ZXN5 | 266:356 | *S. aureus* |
| CBD34 | PlyPMBT8_SH35 | A0A4Y6EB79 | 202:290 | *Staphylococcus hyicus* |
| CBD35 | Ply2638A_SH35 | Q4ZD58 | 398:486 | *S. aureus* |
| CBD36 | UCP1_SH35 | A0A2H4JC36 | 400:480 | Non-annotated host |
| CBD37 | IME1323_01_CBD | A0A1W6JNV3 | 342:440 | *Staphylococcus caprae* |
| CBD38 | PlyIMEP5_CBD | A0A1D6Z273 | 354:456 | *S. aureus* |
| CBD39 | UCP2_SH35 | A0A2H4JBD0 | 368:462 | Non-annotated host |
| CBD40 | SPbeta_CBD | A0A0N9SGX6 | 382:500 | *S. epidermidis* |
| CBD41 | IPLA7_CBD | I6TG35 | 361:460 | *S. epidermidis* |
| CBD42 | Romulus_CBD | M4I0H0 | 173:305 | *S. aureus* |
| CBD43 | phiETA_CBD | Q93R23 | 198:299 | *S. aureus* |
| CBD44 | phiV83_CBD | Q9MBN4 | 136:251 | *S. aureus* |
| CBD45 | phi575_CBD | A0A1Q1PVY4 | 179:289 | *Staphylococcus sciuri* |
| CBD47 | Andhra_CBD | A0A1S6L1H5 | 178:299 | *S. epidermidis* |
| CBD48 | IME1365_01_CBD | A0A1W6JQL2 | 424:569 | Non-annotated host |
| CBD49 | UCP3_PG1 | A0A2H4J236 | 152:293 | Non-annotated host |
| CBD50 | f2b1_PG1 | A0A6C0RTQ9 | 176:318 | *S. aureus* |

### 2.6. Protein expression and purification

The eGFP-CBD fusion proteins were expressed from an *E. coli* BL21(DE3) host transformed with the corresponding expression vector. One hundred mL of terrific broth (TB) were inoculated with an overnight culture of the producing strain and then incubated until reaching OD_600_ ≈ 0.5. The cultures were then induced with 1 mM isopropyl β-D-thiogalactopyranoside (IPTG) and further incubated overnight (∼16 h) at 16 °C with shaking. Finally, the cultures were centrifuged (7 000 × g, 5 min, 4 °C), the supernatants discarded and the pellets resuspended in 1 mL of binding buffer (20 mM sodium phosphate buffer pH 7.4, 300 mM NaCl, 20 mM imidazole) containing 10 µg/mL DNase I and 0.1 mM phenylmethylsulfonyl fluoride (PMSF).

These cell suspensions were disrupted by applying three freeze-thawing cycles plus sonication at 40% amplitude (5 s on/off pulses for 2 min). Samples were centrifuged again to remove cell debris, and the supernatants were transferred to new tubes to which 200 µl of Ni-coated agarose beads (HisPur™ Ni-NTA Superflow Agarose) were added. Then the mixtures were incubated overnight (∼16 h) in an end-over-end shaker at 4 °C. After centrifugation (700 × g, 2 min, 4 °C), the supernatant was removed and the pellet was washed once with 1.5 mL binding buffer and twice with 1.5 mL wash buffer (20 mM sodium phosphate buffer pH 7.4, 300 mM NaCl, 50 mM imidazole). Finally, the purified proteins were recovered by resuspending the beads in 500 µL elution buffer (20 mM sodium phosphate buffer pH 7.4, 300 mM NaCl, 500 mM imidazole) and recovering the supernatant after a new centrifugation. Buffer exchange was performed using a desalting plate (Zeba™ Spin Desalting Columns) according to the instructions of the manufacturer with PBS (137 mM NaCl, 2.7 mM KCl, 10 mM Na_2_HPO_4_, 1.8 mM KH_2_PO_4_, pH 7.4). Purity was confirmed by SDS-PAGE (**Supplementary Figure S1**) and protein molar concentrations were calculated using the Bradford assay and the predicted molecular weights of the proteins (available in **Supplementary Figure S1**).

### 2.7. Quantification of CBD bacterial binding

Binding specificity profiles were obtained for the 24 selected eGFP-CBDs against each of the 24 staphylococcal strains (**Table 1**) essentially as described in [14]. Exponential phase cultures (OD_600_ ≈ 0.5) of the strains were centrifuged (10,000 × g, 5 min) and the pellets were washed with PBS. The bacterial suspensions were adjusted to OD_600_ ≈ 1.0 and added to a dark, flat bottom 96-well plate (180 µL per well). Then 20 µL of a 20 µM solution of each eGFP-CBD were added to every well (except the bacterial autofluorescence controls, to which 20 µL of PBS only were added) and the plates were incubated for 10 min at room temperature. After incubation, the plates were centrifuged (1000 × g, 5 min), the supernatants removed and the pellets washed once with PBS and finally resuspended in 200 µL of PBS. Then, 200 µl of 10 µM fluorescein were added to the plate as internal control, as well as positive fluorescence controls for each eGFP-CBD fusion protein (180 µL PBS plus 20 µL of the 20 µM protein stock solution) as the maximum fluorescence value reachable for the protein. Fluorescence was measured in a TECAN Infinite 200 PRO plate reader (TECAN, Männedorf, Switzerland) with excitation/emission wavelengths of 485 nm and 530 nm, respectively. Fluorescence measurements were acquired for three biological replicates and averaging the values of multiple positions per well in a 3×3, filled circle geometry (thus with a total of 5 measurements per well).

### 2.8. Binding data analysis

Fluorescence measurements were corrected for comparability between proteins by applying a correction factor F_max_/F_prot_ in which F_max_ is the maximum fluorescence recorded among all GFP-CBD constructs and F_prot_ is the fluorescence of each eGFP-CBD at 2 µM. The corresponding blank (the bacterial substrate autofluorescence) was subtracted from each fluorescence data point and the three replicates are presented here as mean ± standard deviation. Specificity profiles (*e.g.* in **Figure 5**) were produced by standardizing the fluorescence values in a 0-1 scale dividing by the highest value reached for each eGFP-CBD in combination with a bacterium. All calculations were performed in R.

Further analyses were performed to obtain insights from the collected data. Particularly, Pearson correlation plots of the binding data were obtained using the “Hmisc” and “corrplot” packages [33,34], considering a significance level of 0.05. Data representation was done with “ggplot2” [35].

## 3. RESULTS

### 3.1. Most staphylococcal CBDs belong to SH3b superfamily

The CBDs from endolysins in PhaLP annotated with a *Staphylococcus* host were extracted and finely delineated by reference to a set of predicted three-dimensional structures. Based on these delineated sequences, a phylogeny of staphylococcal CBDs was built (**Figure 1**). The tree also shows the selected examples for 3D modelling (green/red), and those instances selected for further experimental testing (red) and further annotations (see Materials and Methods and the legend to the figure).

In the course of probing the 3D structures of the 42 representative examples, it was found that all of the models but two (the ones predicted to be *PG_binding_1* [PF01471] domains) displayed structures containing only β-sheets, which aligned fairly well (RMSD between 0.5 and 2.5 according to DALI pairwise comparisons) with a *bona fide SH3_5* (PF08460) CBD, extracted from the crystal structure of lysostaphin (PDB 5LEO).

However, not all of them were predicted, according to Pfam, as part of the *SH3_5* family, which led us to believe that they were different, still undescribed SH3b-like folds. Therefore, we performed an all-against-all DALI structural alignment of the 3D models of the SH3b-like CBDs (**Supplementary File S1**, **Figure 2**) to verify their structural taxonomy. According to the Z-score pairwise similarity matrix (**Figure 2A, Supplementary Table S3**), the structures group together in two large clusters, hereby named *SH3b_P* and *SH3b_T*. The former is in turn divided in two sub-clusters, *SH3b_P1* and *SH3b_P2*. All the representatives in *SH3b_P1* are predicted to be *SH3_5* domains, whereas *SH3b_T* corresponds to the recently described [14] family included as Pfam entry PF24246, and *SH3b_P2* is not consistently associated to any established family in Pfam. When comparing the 3D models from each defined group, *SH3b_P1* and *SH3b_P2* seem to share similarly predicted folds (**Figure 2B**), with the same topology (**Figure 2C**). On the contrary, whereas *SH3b_T* CBDs share the barrel-like β-sheet organization of SH3b folds, their predicted shape is clearly different to *SH3b_P*. Specifically, the topological representation shows that *SH3b_P* and *SH3b_T* differ substantially in the configuration of their N- and C-terminal β-sheets, sharing the structure of the central three-β-sheets core only. An architectural difference can be added too: while *SH3b_P* representatives occurred exclusively as single repeats in our set of endolysins, *SH3b_T* ones appeared either as single or two repeats.

Based on these results, the full set of CBDs were structurally classified within the four possible structural groups or CBD families (*SH3b_P1*, *SH3b_P2*, *SH3b_T* and *PG_binding_1*), depending on the type of folds found in the structural models of the examples within their phylogenetic cluster (**Figure 1**).

### 3.2. Most staphylococcal endolysins are bicatalytic and some are interrupted by introns

The possible architectures (combinations of domains within an endolysin) in which these CBDs appear are schematically depicted in **Figure 3A**. The majority of architectures (∼80% of our examples) are bicatalytic, bearing two different predicted catalytic domains at the N-terminal end (one *CHAP* domain –or a peptidase domain in only six cases– and an *Amidase_2* or *Amidase_3* domain) plus an SH3b CBD at the C-terminus. The role of the central amidase domain still remains controversial, since it has been suggested that it acts as an auxiliary binding domain rather than as a proper catalytic domain, at least in endolysins LysSA12 and LysSA97 [36]. The different functional domains are connected to each other by relatively long linkers (roughly between 14 and 25 amino acids long) according to our 3D structure predictions, as depicted in **Figure 3C**. These long linkers may play a relevant role in the activity of the endolysins, either by allowing the autonomous function of each domain or by ensuring the right cooperation between domains, or both. While some entries in our dataset are apparently bimodular endolysins with just one EAD and one CBD, many of these were found to be a part of full bicatalytic endolysins interrupted in the phage genome by the insertion of another gene (such as a predicted HNH homing endonuclease) or a short nucleotide sequence (examples are provided in **Supplementary Figure S2**). These architectures are counted as bicatalytic in **Figure 3A** whenever the insertion was found to contain an endonuclease-coding gene or a virulence factor (namely staphylokinase in A0A345AQJ3). This is because, in these cases, such insertions are assumed to behave as mobile introns, which are self-spliced after transcription to express the full-length, bicatalytic product [37–39]. Our collection of endolysin sequences includes examples of insertions at different sites (between the *CHAP* domain and the amidase domain, or between the amidase domain and the CBD), some of them containing what seems like a functional intron and others with what seem just the remnants of a prior insertion (*e.g.* A0A6F8Z5Y5 contains a four-nucleotide spacer between two endolysin genes), and even an example (A0A060AB53) with two different insertion events. A schematic recreation of the possible genetic history of such intron-containing, *SH3b_P1*-bearing endolysins is shown in **Figure 3B**. No *SH3b_P1* strictly bimodular representatives (*i.e.*, without the remnants of a *CHAP* domain upstream in the genome) were found in our dataset, which supports the notion that *SH3b_P1* CBDs have evolved to specifically work in a bicatalytic context.

**Figure 3.**
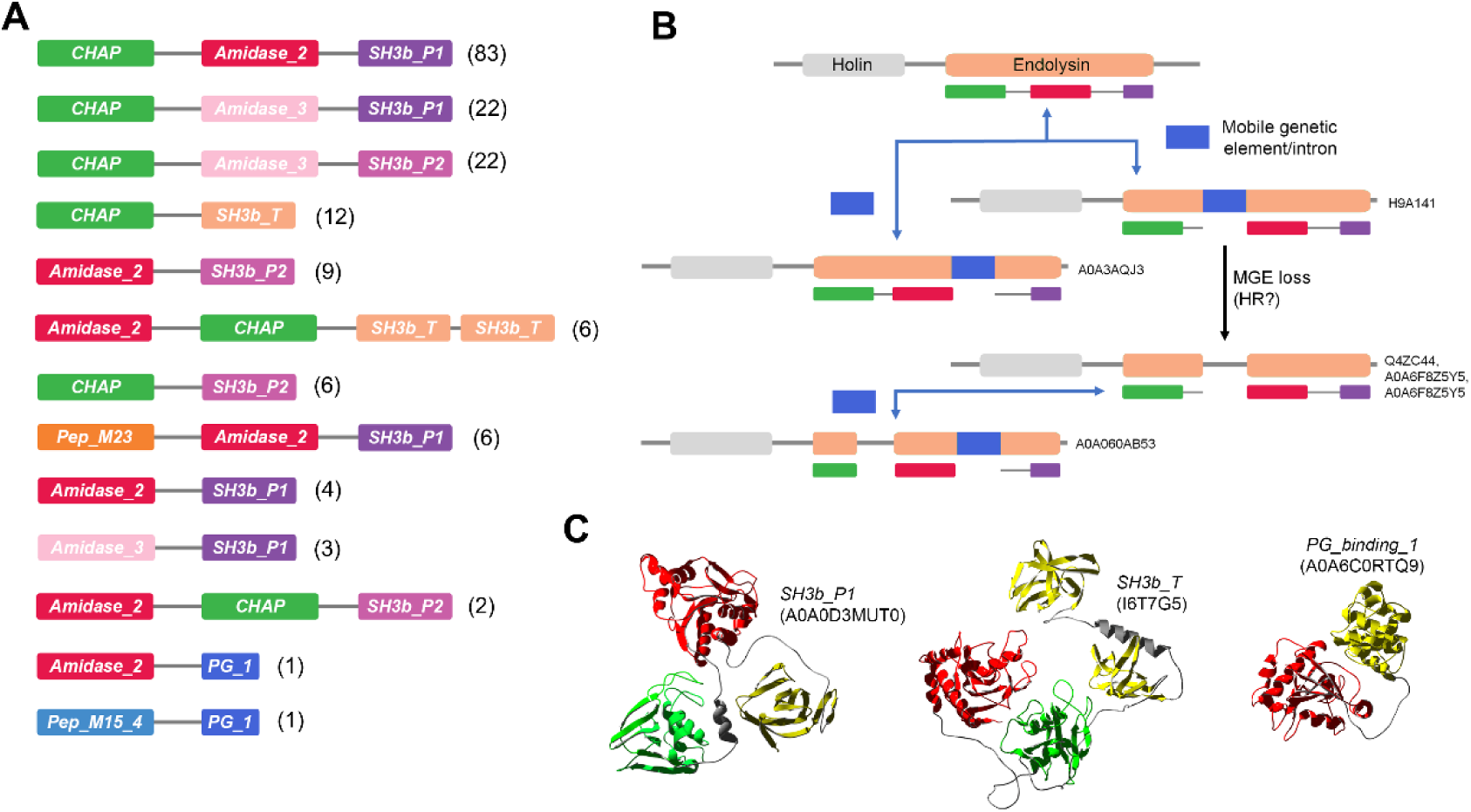
Architectures and genomic arrangements found among the staphylococcal phage endolysins. (A) Schematic representation of the different kinds of architectures predicted in the dataset. Numbers at the right indicate the count of examples of each architecture. Those bimodular architectures for which a putative intron (*i.e.*, an insertion containing a gene coding for an endonuclease or a virulence factor) was detected in the genome were counted as bicatalytic. (B) Proposed routes for the generation of the diverse genomic structure variants observed in our database for *SH3b_P1*-containing endolysins, including the UniProt accession of endolysins for which the depicted genomic arrangements were observed. The *CHAP*, *Amidase_2*, and CBD domains are colored green, red, purple, respectively. (3) Structural models of three architectural examples. *CHAP* domains are shown in green, *Amidase_2* domains in red and CBDs in yellow (with the corresponding CBD type annotated on top of the model).

Purely bimodular architectures, without the presence of an upstream insertion scar, do happen both in endolysins with *SH3b_P2* and *SH3b_T* CBDs, some with a *CHAP* EAD (in *SH3b_P2* and *SH3b_T* examples) and others with an *Amidase_2* or *Amidase_3* EAD (only with *SH3b_P2*). Some *SH3b_T* endolysins also contain two distinct EADs, *Amidase_2* and *CHAP*, but, interestingly, they occur in a different order than when accompanied by an *SH3b_P1* CBD, with the *CHAP* domain in the center of the protein. In addition, the CBD of the latter bicatalytic endolysins (*Amidase_2*:*CHAP*:*SH3b_T*) is always composed by two *SH3b_T* repeats, unlike the case of the bimodular ones (*CHAP*:SH3b_T), in which just a single *SH3b_T* repeat occurs.

### 3.3. Functional characterization of the anti-staphylococcal CBDs

As a next step, a selected set of 24 CBDs were functionally characterized by creating eGFP-CBD fusions and testing their binding against a set of 24 staphylococcal strains. All fusion proteins were created, expressed and purified using immobilized metal affinity chromatography. A number of 24 strains covering 10 staphylococcal species was co-incubated with the fluorescently labelled CBDs and after washing the fluorescence was measured as a metric for binding specificity. A high variability was observed across the specificity profiles (**Supplementary Figure S4**). Specific trends can be extracted from the correlation analyses depicted in **Figure 4**, in which either (A) the specificity profiles of the different CBDs are pairwise compared, thus generating clusters of CBDs with similar binding patterns; or (B) the co-occurrence of binding to each pair of strains is examined, thus revealing how likely it is for any given CBD in our collection to be a good binder for the two strains in each pair simultaneously.

**Figure 4.**
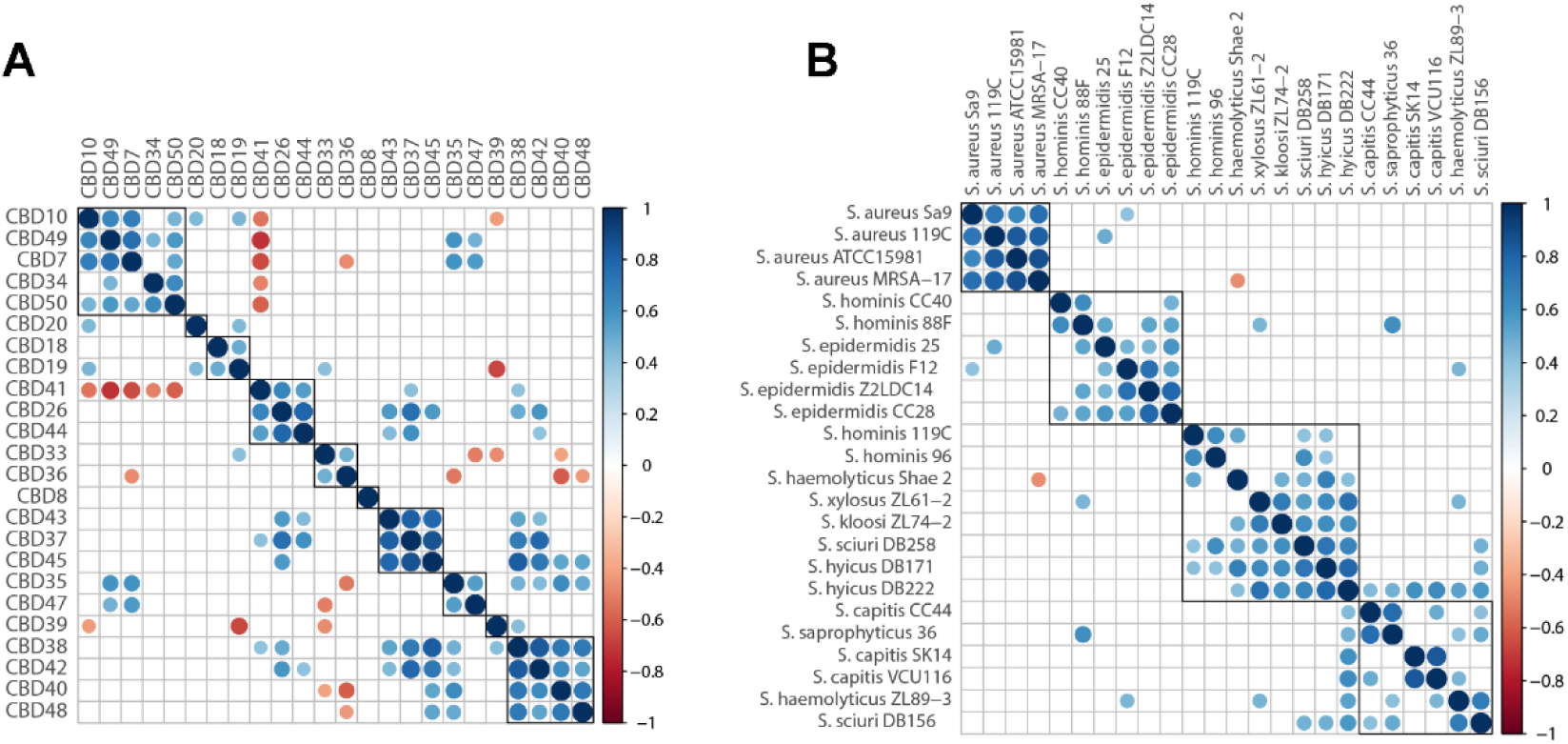
Correlation plots based on the specificity profiles of the CBDs (**Supplementary Figure S4**). The Pearson correlation was calculated on the fluorescence measurements standardized to a 0-1 scale by the minimum and maximum fluorescence values achieved for each eGFP-CBD protein. The different elements (CBDs in [A] or staphylococcal strains in [B]) were ordered by hierarchical clustering based on the correlation coefficients. Correlation coefficients are only displayed when *p*-value ≤ 0.05.

The CBD clusters derived from **Figure 4A** and expanded in the binding profiles of **Figure 5** show certain characteristic binding patterns among the tested CBDs: (i) functional cluster A groups CBDs whose binding is biased towards *S. aureus*/*S. epidermidis*/*S. hominis*; (ii) CBD20, which is a singleton, binds *S. epidermidis*/*S. hominis*/*S. haemolyticus* but not *S. aureus;* (iii) cluster C contains the CBDs with the broadest range; (iv) the representatives in cluster D bind rather specifically *S. haemolyticus* Shae 2 plus others (such as *S. hominis* 119C or *S. kloosi*); (v) the CBDs of cluster E can be better defined by binding non-aureus staphylococci; (vi) the singleton CBD8 binds *S. aureus* and *S. hominis* plus other varied staphylococci (*S. haemolyticus*, *S. sciuri*, *S. hyicus*, *S. saprophyticus*, etc.) but not *S. epidermidis*; (vii) cluster G shows specificity towards *S. hominis* plus *S. haemolyticus* Shae 2; (viii) in cluster H we find CBD35, which shows a specificity profile akin to cluster A (*S. aureus*/*S. epidermidis*/*S. hominis*), together with CBD47, which seems *S. aureus*-specific – however the maximum RFU achieved for eGFP-CBD47 is very low compared to the ones obtained for the other proteins, which points out to inferior binding or defective folding by this particular protein; (ix) CBD39 is, again, a singleton, although its binding profile reminds to that of CBD8 (*i.e.*, broad range excluding *S. epidermidis*) but with higher binding to *S. capitis*; (x) the final cluster is broad range minus *S. capitis*, *S. hominis* 88F and *S. saprophyticus*.

**Figure 5.**
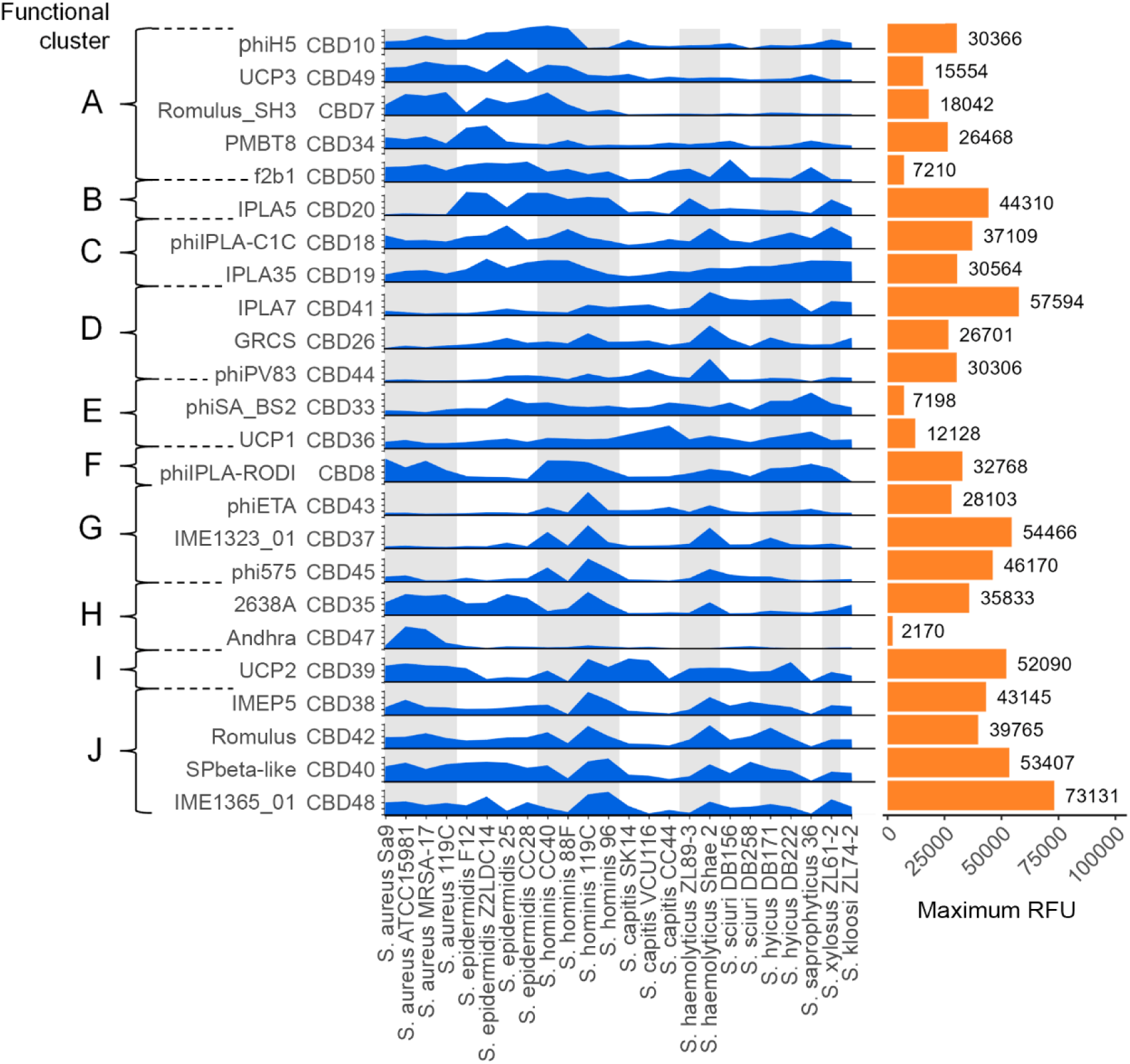
Normalized binding profiles obtained for the experimentally tested eGFP-CBD fusions against a collection of staphylococcal strains. Each CBD is labeled with the name of its original phage (phiH5, GRCS, UCP1, etc.) and its ID (CBD10, CBD26, CBD36, etc.). Blue ridges indicate the measured fluorescence intensity relative to the maximum value for each CBD in a 0-1 scale (Y-axis) and for each tested strain (X-axis). The sets of strains belonging to the same species are highlighted with intercalating light gray and white background shades. Bar plots at the right depict the maximum RFU achieved for each CBD. CBDs are arranged according to the hierarchy in Figure 4A.

**Figure 4B** provides another general reading of these results, focusing on the correlation between strains. The main conclusion from this plot is that the binding measurements to *S. aureus* strains (at least the ones in our study) are highly correlated. In other words, we may say that any CBD –in our test group– that binds an *S. aureus* strain will also bind any other *S. aureus* strain tested. Similarly, binding of *S. epidermidis* strains are also relatively correlated with the other *S. epidermidis* strains. The self-correlation within species is nonetheless not generalized in our dataset. For example, the *S. haemolyticus* and *S. hominis* strains appear split between two different correlation clusters, which means that binding a certain *S. haemolyticus* or *S. hominis* strain does not imply necessarily binding any representative of the species.

Finally, we connected the experimental, functional data with the previously established structural knowledge. **Figure 6** shows the classification of the functionally characterized CBDs in three different categories: annotated host of the phage, functional cluster, and phylogenetic cluster. The first feature was obtained from the metadata of each entry upon retrieval from PhaLP, the second one was calculated from the experimental data as shown in **Supplementary Figure S4** and the third one was obtained from the phylogenetic tree in **Figure 1**. This latter one represents the general structural features of each CBD in the form of a simple classification based on its primary amino acid sequence. One main conclusion can be derived from the general picture of the connections in **Figure 6**: neither the original bacterial host nor the phylogenetic cluster are good predictors for the observed specificity profile. This is supported by the fact that each functional class is connected to many different host classes or phylogenetic classes, which may be taken as an indicator of intense horizontal gene transfer in the staphylococcal bacteria-phage interface. In other words, there are no clearly observable correlations between the classes presented in **Figure 6**.

**Figure 6.**
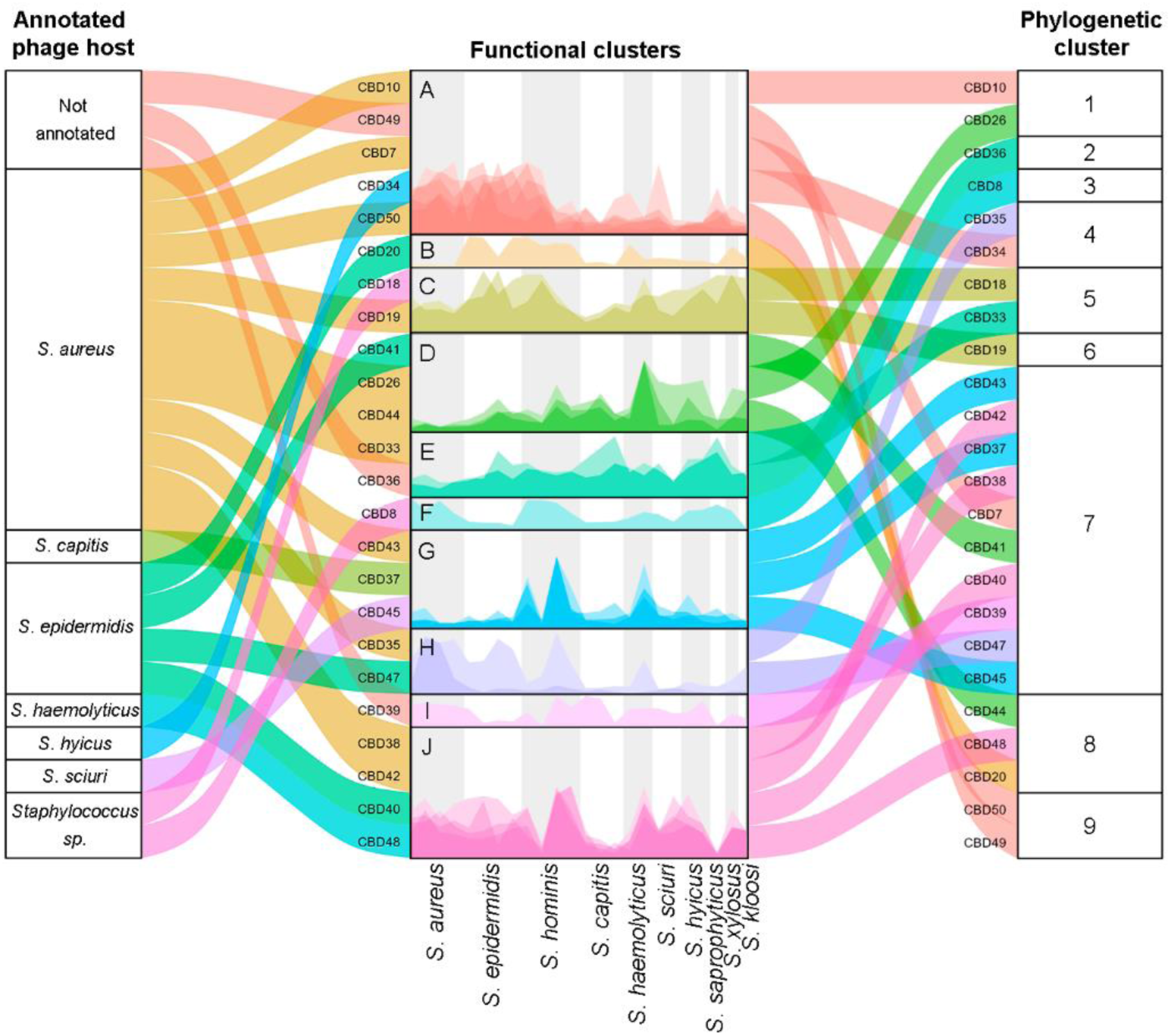
Relationships between the functional clusters of CBDs, the annotated host of the source phage and the phylogenetic cluster as shown in Figure 1. Each stripe depicts a single CBD. In the connections between annotated host and functional cluster, different colors represent different bacterial hosts. In the functional cluster strata and the connections between them and the phylogenetic clusters, color codes refer to the different functional clusters. The specificity profiles from Figure 5 are shown in the central column, overlaying all profiles for each particular cluster. Y-axis is not scaled but rather re-dimensioned in each stratum to fit its height. The different species of the target bacteria for the specificity profiles are signaled by grey-white alternating vertical shades and annotated at the bottom of the central column.

## 4. DISCUSSION

Our observations show that an overwhelming majority of staphylococcal endolysins contains CBDs based on SH3b folds. These CBDs do not only belong to known SH3b families [17], but also to at least two more families of undescribed CBDs (*SH3b_T* and *SH3b_P2*, a sub-family of the broader *SH3b_P* family, which contains the well-known *SH3_5*, herewith named *SH3b_P1*). Furthermore, a relevant amount of staphylococcal endolysins are interrupted by DNA sequences likely containing self-splicing introns. The presence of introns further complicates the proper annotation of these phage genomes. Many of the endolysins currently annotated as bimodular in the public repositories actually have three modules when expressed, which should be taken into account when analyzing newly sequenced staphylococcal genomes. The genome of phage K (AY176327.1) can be taken as an example in which the endolysin is correctly annotated, including its inserted intron. Of note, we only found these insertions in genes coding for endolysins with an *SH3b_P1* CBD, which, in turn, are the preferred ones in phages from *S. aureus* hosts. Previous literature already hinted that intron insertions are a typical feature of twortlikeviruses infecting *S. aureus* [39], and that not all bacterial or phage genomes may be equally susceptible to handle such mobile genetic elements [40]. This explains to a certain extent the absence of introns in the endolysin genes that have evolved to accommodate CBDs other than *SH3b_P1*. Thus, the conditions for intron acquisition may have occurred after the evolutionary divergence of *SH3b_P1* and *SH3b_P2* (and this divergence was after the one of *SH3b_P* and *SH3b_T*). The possible function that these introns may fulfill in the phage context is still not sufficiently elucidated in current literature. It has been proposed that introns may be beneficial by providing noncoding targets for the recombination of exons (as shown earlier for T4 virus [41]) or that they may allow the synthesis of several proteins from the same gene via incomplete or alternative splicing [37]. While the former reason fits well with the dynamic and mosaic nature of phage genomes, the latter can also be accepted since many phage endolysins are now known to encode secondary ribosome binding sites (RBSs) to express additional, shorter monomers that play a role in facilitating the activity of the endolysin [42]. Incidentally, an attempt at predicting secondary RBSs in the representative set of 42 endolysins was inconclusive (**Supplementary Figure S3**), as anyway shown before for staphylococcal endolysins [42], even though there is at least one experimentally proven example, the endolysin of phage 2638A, with a functional RBS between its two EADs (*Peptidase_M23* and *Amidase_2*) [43]. Another remarkable observation is the occurrence of *SH3b_T* domains as two repeats only when in an architecture bearing two EADs. A possible explanation would be that, in this case, the two EADs synergistically cooperate to render a more efficient enzyme, which then may be required to bind very tightly to the target cell wall to avoid lysis of neighboring and potentially new host cells. Thus, they may have acquired a second *SH3b_T* repeat to increase ligand affinity. This is supported by the higher maximum RFU achieved by CBD20 (44310) and CBD48 (73131) as compared with CBD44 (30306) (**Figure 5**), the latter coming from a *CHAP:SH3b_T* architecture and the former two from an *Amidase_2:CHAP:SH3b_T:SH3b_T* one.

As a general conclusion so far, we may say that certain domain configurations are preferred by specific phylogenetic clusters of CBDs (**Figure 1**), or, to put it differently, the particular primary sequence of a CBD correlates with certain architectural features of the endolysin in which it is found. Some examples of this kind of correlations are: (i) *SH3b_P1* was never found within a bicatalytic architecture with the order Amidase:*CHAP*:CBD; (ii) the latter bicatalytic architecture was always accompanied by two *SH3b_T* repeats instead of just a single one, or a single *SH3b_P2*; (iii) potential self-splicing introns were found only in architectures with CBDs of the *SH3b_P1* type. These domain correlations suggest a functional counterpart: that a strong relationship or even cooperation between the different modules occurs towards fulfilling the physiological function of the protein, rather than just a merely autonomous work of each domain. From an evolutionary point-of-view, we may say then that some functional constraints operate in which certain CBD sequences are positively selected only in the company of certain combinations of EAD(s). This aligns with the notion that interactions mediated by SH3 domains are strongly influenced by the “protein context” in which they are integrated [44].

Focusing on the experimental characterization of the binding profiles of our selected CBDs, the most salient feature was the high self-correlation of binding to *S. aureus* strains, while no similarly self-correlated binding clusters were obtained for other staphylococcal species except, perhaps, *S. epidermidis* (**Figure 4B**). This suggests that the ligand(s) recognized at the surface of *S. aureus* cells are conserved across all the tested *S. aureus* strains. Since it is known that at least some *SH3b_P1* CBDs bind directly to the peptidoglycan, including its peptide cross-link [45,46], it is easy to signal a shared specific trait of *S. aureus* that is not widespread in other staphylococci and could serve as ligand: the pentaglycine cross-link. Other ligands may, however, be also involved in this phenomenon. This, in turn, and with all due cautions given the small population size, points out to a more variable configuration of surface ligands targeted by the CBDs in other species than *S. aureus* or *S. epidermidis*.

Judging by the aforementioned results, and as specially seen in **Figure 5**, it may be concluded that species-level or strain-level specificity is rarely achieved by the tested staphylococcal CBDs, since most of them display overarching specificities that extend across many staphylococcal species. This contrasts with the exquisite specificity previously described for *Listeria* or *Streptococcus pneumoniae* phage endolysins [47,48]. A possible explanation for this is the nature of the ligands that the respective staphylococcal CBDs bind. While the *Listeria* and S*. pneumoniae* CBDs seem to bind substituents at the teichoic acids, which may greatly vary at the serovar/strain or species level, at least some of the SH3b CBDs in staphylococcal CBDs are known to bind different parts of the peptidoglycan, which is a more conserved structure.

However, given our experimental approach, a couple of clarifications must be made. Firstly, the observed specificities could depend on the concentration of the CBD. Our assays were performed at 2 µM, which is in the higher end but still within the usual dosage range of endolysins when applied exogenously as antimicrobials against Gram-positive bacteria. It is thus a relevant concentration from an applied point of view, although not necessarily from a physiological point of view (*i.e.*, to account for a realistic setting in the release step of the phage infection cycle). Likewise, our procedure for assessing CBD binding relies on the interaction of the eGFP-CBDs with the bacterial surface from without the cell. This contrasts with the actual mechanism occurring during the phage infection cycle in which endolysins (and thus CBDs) are released into the cell wall from within the cell. When this release occurs, it is normally associated with the collapse of the proton motive force, which causes a modification of the chemical environment at the cell wall known to significantly impact the behavior of endolysins [49]. These physiological conditions are not taken into account in our experimental setup. Thus, our observations so far may be more relevant towards the applied development of CBDs rather than to their natural role in the endolysin –which may be a limitation of our approach but does not completely preclude us to derive evolutionary consequences for the phage-bacteria system, especially when clear correlations at the taxonomical level are observed (*e.g.*, the case of *S. aureus*).

This taken into account, and considering the lack of meaningful correlations between structure, host identify and specificity profile as presented in **Figure 6**, we propose two general evolutionary conclusions from this work:

(i) The specificity profile itself does not seem a major evolutionary constraint in the context of the staphylococci-phage system, since there are no clear “preferences” of any host for a singular CBD functional profile. While we cannot claim this is a general trend among all staphylococci given the limited nature of our study, it is *e.g.* clearly denoted by the CBDs derived from phages with an *S. aureus* annotated host. In this case, we see (**Figure 6**) that these CBDs are represented in seven out of ten functional clusters, and some of them (clusters D and G) are not even particularly optimized for *S. aureus* binding specificity, at least from without the cell. These observations are somewhat against the common notion that an endolysin would be most specific towards the bacterium that acts as a host for its phage. Two theoretical options may be taken from this conclusion: (a) either we maintain the notion that CBD specificity is a feature evolved in endolysins and we are not observing it here due to our experimental setting, or, conversely, (b) we reject the CBD host specificity as a preferentially evolved trait within the phage. It is our point-of-view that it is not immediately obvious why bacterial specificity should be, in general, an evolutionarily selected trait for phage endolysins. This is because at the time of the release of the mature virions, the phage has already found a suitable host. Finding such a host justifies the specificity mechanisms associated to virion parts such as the receptor-binding proteins, but it is not well understood why the endolysin should also carry specificity determinants. Thus it can be construed that specificity itself may not be an evolved trait but rather a consequence of evolutionarily selecting CBDs that bind essential or conserved ligands (as is the case of peptidoglycan for at least some staphylococcal CBDs or the phosphocholine residues at the teichoic acids of *S. pneumoniae*) to withstand easy resistance development by the infected bacterium.
(ii) A comparable situation occurs in the connection between functional and phylogenetic clusters (**Figure 6**): each functional cluster comprises CBDs from different phylogenetic clusters, with the only exception of cluster G – in turn, the least broad-range one, whose representatives belong to the same phylogenetic group (cluster 7, which contains all *SH3b_P2* representatives). What similar specificity profiles signal is fundamentally something about access to the targeted ligand at the bacterium, *i.e.*, when certain strains are bound by the same CBD this means that they contain ligands that can be accessed and recognized by the same CBD. And, subsequently, if a group of CBDs display the same pattern of specificity this means that they most likely recognize the same type of ligand present in the same strains. However, the lack of correlation between specificity profiles and phylogenetic groups suggests that recognizing the same ligand can be achieved through different structural solutions (*e.g.*, cluster A contains *SH3b_P1*, *SH3b_P2* and *PG_binding_1* representatives; cluster D contains examples from *SH3b_P1*, *SH3b_P2* and *SH3b_T* families) or, at least, that the specificity is ultimately determined by certain key amino acids [14,46] rather than by the altogether primary sequence.

Although more sophisticated strategies were attempted at discovering meaningful structure-function connections such as single-position specificity determinants (*e.g.*, by applying machine learning algorithms treating each sequence position as a feature to uncover those positions with a greater involvement in predicting the functional clusters) no informative results or insights were obtained. This is related to the fact that the experimentally tested subset is too small to detect the interplay of likely multiple variable amino acids that ultimately govern the observed binding specificity. And this should also be a major conclusion from this work: although the staphylococcal CBDs have evolved very similar structures (*i.e.*, most of them with a predicted SH3b-like fold), rather subtle differences in sequence render relatively radical changes in observed specificity. As such, staphylococcal CBDs have evolved radically different to highly specific CBDs acquired by *Listeria* or *S. pneumoniae* phages.

## 5. CONCLUSION

With this work, we have probed the structure-function relationships of CBDs derived from staphylococcal phage endolysins. We have shown that most of the known diversity of staphylococcal CBDs adopts an SH3b fold, although in different related families and sub-families. We have also demonstrated the possibility to systemically obtain experimental specificity profiles by generating eGFP-CBD fusions and testing them in a semi-high throughput manner against a panel of test strains. This information already poses a valuable, direct addition towards the engineering of exogenously applied endolysins, but it can also be used as a base for the discussion of more fundamental insights. From an evolutionary standpoint, the hereby presented data challenge the notion of CBD specificity as a primarily selected trait in endolysins, at least within the staphylococcal context. And, from a structural point of view, our results suggest that (i) while a dynamic exchange of modules between phages/hosts is possible, CBDs and EADs are evolutionarily fixed in certain preferred combinations which hint at extensive cooperation between both types of domains within the endolysin; and (ii) the main specificity determinant for staphylococcal CBDs is likely not the general structure, nor its domain family, but rather key amino acid residues that can only be uncovered by in-depth biochemical characterizations of the receptor-ligand contact or, alternatively, by deriving sufficiently informative experimental data sets.

## Data and resource availability

### Funding sources

This work was supported by the Research Foundation—Flanders (FWO) under Research project G066919N and PhD SB fellowship 1S38519N to B.C. R.V. was supported by a postdoctoral fellowship of the “Bijzonder Onderzoeksfonds” (BOF) Ghent University (01P10022).

### Conflicts of interest

B.C. and Y.B. are co-founders and employee (B.C.) or scientific advisor (Y.B.) of Obulytix. D.G. is currently employed by Telum Therapeutics. R.V. has provided scientific consulting services to Obulytix.

### Author Contributions

R.V.: conceptualization, data curation, formal analysis, investigation, methodology, visualization, writing – original draft, writing – review and editing; D.G.: conceptualization, data curation, investigation, methodology, supervision, writing – review and editing; B.C.: data curation, formal analysis, writing – review and editing; Z.D.: investigation; Y.B.: conceptualization, project administration, resources, supervision, writing – review and editing.

## REFERENCES

[1] World Health Organization;, 2019 Antibacterial agents in clinical development: an analysis of the antibacterial clinical development pipeline, World Health Organization (2019). https://www.who.int/publications/i/item/9789240000193.

[2] U. Theuretzbacher, K. Outterson, A. Engel, A. Karlen, The global preclinical antibacterial pipeline, Nat Rev Microbiol 18 (2020) 275–285. 10.1038/s41579-019-0288-0.

[3] Vázquez R, García E, García P, Phage lysins for fighting bacterial respiratory infections: a new generation of antimicrobials, Front Immunol 9 (2018) 2252. 10.3389/fimmu.2018.02252.

[4] D. Gutierrez, L. Fernandez, A. Rodriguez, P. Garcia, Are phage lytic proteins the secret weapon to kill *Staphylococcus aureus*?, mBio 9 (2018). 10.1128/mBio.01923-17.

[5] D. Dams, Y. Briers, Enzybiotics: enzyme-based antibacterials as therapeutics, Adv Exp Med Biol 1148 (2019) 233–253. 10.1007/978-981-13-7709-9_11.

[6] M. Schmelcher, M.J. Loessner, Bacteriophage endolysins — extending their application to tissues and the bloodstream, Current Opinion in Biotechnology 68 (2021) 51–59. 10.1016/j.copbio.2020.09.012.

[7] R. Vázquez, E. García, P. García, Sequence-Function Relationships in Phage-Encoded Bacterial Cell Wall Lytic Enzymes and Their Implications for Phage-Derived Product Design, J Virol 95 (2021) e0032121. 10.1128/JVI.00321-21.

[8] B. Criel, S. Taelman, W. Van Criekinge, M. Stock, Y. Briers, PhaLP: A Database for the Study of Phage Lytic Proteins and Their Evolution, Viruses 13 (2021). 10.3390/v13071240.

[9] D. Botstein, A theory of modular evolution for bacteriophages, Ann N Y Acad Sci 354 (1980) 484–490. 10.1111/j.1749-6632.1980.tb27987.x.

[10] B.J. Smug, K. Szczepaniak, E.P.C. Rocha, S. Dunin-Horkawicz, R.J. Mostowy, Ongoing shuffling of protein fragments diversifies core viral functions linked to interactions with bacterial hosts, Nat Commun 14 (2023) 7460. 10.1038/s41467-023-43236-9.

[11] E. Diaz, R. Lopez, J.L. Garcia, Chimeric pneumococcal cell wall lytic enzymes reveal important physiological and evolutionary traits, J Biol Chem 266 (1991) 5464– 71.

[12] H. Gerstmans, B. Criel, Y. Briers, Synthetic biology of modular endolysins, Biotechnol Adv 36 (2018) 624–640. 10.1016/j.biotechadv.2017.12.009.

[13] H. Gerstmans, D. Grimon, D. Gutierrez, C. Lood, A. Rodriguez, V. van Noort, J. Lammertyn, R. Lavigne, Y. Briers, A VersaTile-driven platform for rapid hit-to-lead development of engineered lysins, Sci Adv 6 (2020) eaaz1136. 10.1126/sciadv.aaz1136.

[14] R. Vázquez, D. Gutiérrez, D. Grimon, L. Fernández, P. García, A. Rodríguez, Y. Briers, The New SH3b_T Domain Increases the Structural and Functional Variability Among SH3b-Like CBDs from Staphylococcal Phage Endolysins, Probiotics & Antimicro. Prot. (2024). 10.1007/s12602-024-10309-0.

[15] K.S. Ikuta, L.R. Swetschinski, G.R. Aguilar, F. Sharara, T. Mestrovic, A.P. Gray, N.D. Weaver, E.E. Wool, C. Han, A.G. Hayoon, A. Aali, S.M. Abate, M. Abbasi-Kangevari, Z. Abbasi-Kangevari, S. Abd-Elsalam, G. Abebe, A. Abedi, A.P. Abhari, H. Abidi, R.G. Aboagye, A. Absalan, H.A. Ali, J.M. Acuna, T.D. Adane, I.Y. Addo, O.A. Adegboye, M. Adnan, Q.E.S. Adnani, M.S. Afzal, S. Afzal, Z.B. Aghdam, B.O. Ahinkorah, A. Ahmad, A.R. Ahmad, R. Ahmad, S. Ahmad, S. Ahmad, S. Ahmadi, A. Ahmed, H. Ahmed, J.Q. Ahmed, T.A. Rashid, M. Ajami, B. Aji, M. Akbarzadeh-Khiavi, C.J. Akunna, H.A. Hamad, F. Alahdab, Z. Al-Aly, M.A. Aldeyab, A.V. Aleman, F.A.N. Alhalaiqa, R.K. Alhassan, B.A. Ali, L. Ali, S.S. Ali, Y. Alimohamadi, V. Alipour, A. Alizadeh, S.M. Aljunid, K. Allel, S. Almustanyir, E.K. Ameyaw, A.M.L. Amit, N. Anandavelane, R. Ancuceanu, C.L. Andrei, T. Andrei, D. Anggraini, A. Ansar, A.E. Anyasodor, J. Arabloo, A.Y. Aravkin, D. Areda, T. Aripov, A.A. Artamonov, J. Arulappan, R.T. Aruleba, M. Asaduzzaman, T. Ashraf, S.S. Athari, D. Atlaw, S. Attia, M. Ausloos, T. Awoke, B.P.A. Quintanilla, T.M. Ayana, S. Azadnajafabad, A.A. Jafari, D.B. B, M. Badar, A.D. Badiye, N. Baghcheghi, S. Bagherieh, A.A. Baig, I. Banerjee, A. Barac, M. Bardhan, F. Barone-Adesi, H.J. Barqawi, A. Barrow, P. Baskaran, S. Basu, A.-M.M. Batiha, N. Bedi, M.A. Belete, U.I. Belgaumi, R.G. Bender, B. Bhandari, D. Bhandari, P. Bhardwaj, S. Bhaskar, K. Bhattacharyya, S. Bhattarai, S. Bitaraf, D. Buonsenso, Z.A. Butt, F.L.C. dos Santos, J. Cai, D. Calina, P. Camargos, L.A. Cámera, R. Cárdenas, M. Cevik, J. Chadwick, J. Charan, A. Chaurasia, P.R. Ching, S.G. Choudhari, E.K. Chowdhury, F.R. Chowdhury, D.-T. Chu, I.S. Chukwu, O. Dadras, F.T. Dagnaw, X. Dai, S. Das, A. Dastiridou, S.A. Debela, F.W. Demisse, S. Demissie, D. Dereje, M. Derese, H.D. Desai, F.N. Dessalegn, S.A.A. Dessalegni, B. Desye, K. Dhaduk, M. Dhimal, S. Dhingra, N. Diao, D. Diaz, S. Djalalinia, M. Dodangeh, D. Dongarwar, B.T. Dora, F. Dorostkar, H.L. Dsouza, E. Dubljanin, S.J. Dunachie, O.C. Durojaiye, H.A. Edinur, H.B. Ejigu, M. Ekholuenetale, T.C. Ekundayo, H. El-Abid, M. Elhadi, M.A. Elmonem, A. Emami, L.E. Bain, D.B. Enyew, R. Erkhembayar, B. Eshrati, F. Etaee, A.F. Fagbamigbe, S. Falahi, A. Fallahzadeh, E.J.A. Faraon, A. Fatehizadeh, G. Fekadu, J.C. Fernandes, A. Ferrari, G. Fetensa, I. Filip, F. Fischer, M. Foroutan, P.A. Gaal, M.A. Gadanya, A.M. Gaidhane, B. Ganesan, M. Gebrehiwot, R. Ghanbari, M.G. Nour, A. Ghashghaee, A. Gholamrezanezhad, A. Gholizadeh, M. Golechha, P. Goleij, D. Golinelli, A. Goodridge, D.A. Gunawardane, Y. Guo, R.D. Gupta, S. Gupta, V.B. Gupta, V.K. Gupta, A. Guta, P. Habibzadeh, A.H. Avval, R. Halwani, A. Hanif, M.A. Hannan, H. Harapan, S. Hassan, H. Hassankhani, K. Hayat, B. Heibati, G. Heidari, M. Heidari, R. Heidari-Soureshjani, C. Herteliu, D.Z. Heyi, K. Hezam, P. Hoogar, N. Horita, M.M. Hossain, M. Hosseinzadeh, M. Hostiuc, S. Hostiuc, S. Hoveidamanesh, J. Huang, S. Hussain, N.R. Hussein, S.E. Ibitoye, O.S. Ilesanmi, I.M. Ilic, M.D. Ilic, M.T. Imam, M. Immurana, L.R. Inbaraj, A. Iradukunda, N.E. Ismail, C.C.D. Iwu, C.J. Iwu, L.M. J, M. Jakovljevic, E. Jamshidi, T. Javaheri, F. Javanmardi, J. Javidnia, S.K. Jayapal, U. Jayarajah, R. Jebai, R.P. Jha, T. Joo, N. Joseph, F. Joukar, J.J. Jozwiak, S.E.O. Kacimi, V. Kadashetti, L.R. Kalankesh, R. Kalhor, V.K. Kamal, H. Kandel, N. Kapoor, S. Karkhah, B.G. Kassa, N.J. Kassebaum, P.D. Katoto, M. Keykhaei, H. Khajuria, A. Khan, I.A. Khan, M. Khan, M.N. Khan, M.A. Khan, M.M. Khatatbeh, M.M. Khater, H.R.K. Kashani, J. Khubchandani, H. Kim, M.S. Kim, R.W. Kimokoti, N. Kissoon, S. Kochhar, F. Kompani, S. Kosen, P.A. Koul, S.L.K. Laxminarayana, F.K. Lopez, K. Krishan, V. Krishnamoorthy, V. Kulkarni, N. Kumar, O.P. Kurmi, A. Kuttik-kattu, H.H. Kyu, D.K. Lal, J. Lám, I. Landires, S. Lasrado, S. Lee, J. Lenzi, S. Lewycka, S. Li, S.S. Lim, W. Liu, R. Lodha, M.J. Loftus, A. Lohiya, L. Lorenzovici, M. Lotfi, A. Mahmoodpoor, M.A. Mahmoud, R. Mahmoudi, A. Majeed, J. Majidpoor, A. Makki, G.A. Mamo, Y. Manla, M. Martorell, C.N. Matei, B. McManigal, E.M. Nasab, R. Mehrotra, A. Melese, O. Mendoza-Cano, R.G. Menezes, A.-F.A. Mentis, G. Micha, I.M. Michalek, A.C.M.G.N. de Sá, N.M. Kostova, S.A. Mir, M. Mirghafourvand, S. Mirmoeeni, E.M. Mirrakhimov, M. Mirza-Aghazadeh-Attari, A.S. Misganaw, A. Misganaw, S. Misra, E. Mohammadi, M. Mohammadi, A. Mohammadian-Hafshejani, S. Mohammed, S. Mohan, M. Mohseni, A.H. Mokdad, S. Momtazmanesh, L. Monasta, C.E. Moore, M. Moradi, M.M. Sarabi, S.D. Morrison, M. Motaghinejad, H.M. Isfahani, A.M. Khaneghah, S.A. Mousavi-Aghdas, S. Mubarik, F. Mulita, G.B.B. Mulu, S.B. Munro, S. Muthupandian, T.S. Nair, A.A. Naqvi, H. Narang, Z.S. Natto, M. Naveed, B.P. Nayak, S. Naz, I. Negoi, S.A. Nejadghaderi, S.N. Kandel, C.H. Ngwa, R.K. Niazi, A.T.N. de Sá, N. Noroozi, H. Nouraei, A. Nowroozi, V. Nuñez-Samudio, J.J. Nutor, C.I. Nzoputam, O.J. Nzoputam, B. Oancea, R.M. Obaidur, V.A. Ojha, A.P. Okekunle, O.C. Okonji, A.T. Olagunju, B.O. Olusanya, A.O. Bali, E. Omer, N. Otstavnov, B. Oumer, M.P. A, J.R. Padubidri, K. Pakshir, T. Palicz, A. Pana, S. Pardhan, J.L. Paredes, U. Parekh, E.-C. Park, S. Park, A. Pathak, R. Paudel, U. Paudel, S. Pawar, H.P. Toroudi, M. Peng, U. Pensato, V.C.F. Pepito, M. Pereira, M.F.P. Peres, N. Perico, I.-R. Petcu, Z.Z. Piracha, I. Podder, N. Pokhrel, R. Poluru, M.J. Postma, N. Pourtaheri, A. Prashant, I. Qattea, M. Rabiee, N. Rabiee, A. Radfar, S. Raeghi, S. Rafiei, P.R. Raghav, L. Rahbarnia, V. Rahimi-Movaghar, M. Rahman, M.A. Rahman, A.M. Rahmani, V. Rahmanian, P. Ram, M.M.A.N. Ranjha, S.J. Rao, M.-M. Rashidi, A. Rasul, Z.A. Ratan, S. Rawaf, R. Rawassizadeh, M.S. Razeghinia, E.M.M. Redwan, M.T. Regasa, G. Remuzzi, M.A. Reta, N. Rezaei, A. Rezapour, A. Riad, R.K. Ripon, K.E. Rudd, B. Saddik, S. Sadeghian, U. Saeed, M. Safaei, A. Safary, S.Z. Safi, M. Sahebazzamani, A. Sahebkar, H. Sahoo, S. Salahi, S. Salahi, H. Salari, S. Salehi, H.S. Kafil, A.M. Samy, N. Sanadgol, S. Sankararaman, F. Sanmarchi, B. Sathian, M. Sawhney, G.K. Saya, S. Senthilkumaran, A. Seylani, P.A. Shah, M.A. Shaikh, E. Shaker, M.Z. Shakhmardanov, M.M. Sharew, A. Sharifi-Razavi, P. Sharma, R.A. Sheikhi, A. Sheikhy, P.H. Shetty, M. Shigematsu, J.I. Shin, H. Shirzad-Aski, K.M. Shivakumar, P. Shobeiri, S.A. Shorofi, S. Shrestha, M.M. Sibhat, N.B. Sidemo, M.K. Sikder, L.M.L.R. Silva, J.A. Singh, P. Singh, S. Singh, M.S. Siraj, S.S. Siwal, V.Y. Skryabin, A.A. Skryabina, B. Socea, D.D. Solomon, Y. Song, C.T. Sreeramareddy, M. Suleman, R.S. Abdulkader, S. Sultana, M. Szócska, S.-A. Tabatabaeizadeh, M. Tabish, M. Taheri, E. Taki, K.-K. Tan, S. Tandukar, N.Y. Tat, V.Y. Tat, B.N. Tefera, Y.M. Tefera, G. Temesgen, M.-H. Temsah, S. Tharwat, A. Thiyagarajan, I.I. Tleyjeh, C.E. Troeger, K.K. Umapathi, E. Upadhyay, S.V. Tahbaz, P.R. Valdez, J.V. den Eynde, H.R. van Doorn, S. Vaziri, G.-I. Verras, H. Viswanathan, B. Vo, A. Waris, G.T. Wassie, N.D. Wickramasinghe, S. Yaghoubi, G.A.T.Y. Yahya, S.H.Y. Jabbari, A. Yigit, V. Yiğit, D.K. Yon, N. Yonemoto, M. Zahir, B.A. Zaman, S.B. Zaman, M. Zangiabadian, I. Zare, M.S. Zastrozhin, Z.-J. Zhang, P. Zheng, C. Zhong, M. Zoladl, A. Zumla, S.I. Hay, C. Dolecek, B. Sartorius, C.J.L. Murray, M. Naghavi, Global mortality associated with 33 bacterial pathogens in 2019: a systematic analysis for the Global Burden of Disease Study 2019, The Lancet 400 (2022) 2221–2248. 10.1016/S0140-6736(22)02185-7.

[16] V.G. Fowler Jr., A.F. Das, J. Lipka-Diamond, J.E. Ambler, R. Schuch, R. Pomerantz, C. Cassino, L. Jáuregui-Peredo, G.J. Moran, M.E. Rupp, A.M. Lachiewicz, J.L. Kuti, R.A. Wise, K.S. Kaye, M.J. Zervos, W.G. Nichols, Exebacase in Addition to Standard-of-Care Antibiotics for Staphylococcus aureus Bloodstream Infections and Right-Sided Infective Endocarditis: A Phase 3, Superiority-Design, Placebo-Controlled, Randomized Clinical Trial (DISRUPT), Clinical Infectious Diseases (2024) ciae043. 10.1093/cid/ciae043.

[17] H. Oliveira, M. Sampaio, L.D.R. Melo, O. Dias, W.H. Pope, G.F. Hatfull, J. Azeredo, Staphylococci phages display vast genomic diversity and evolutionary relationships, BMC Genomics 20 (2019) 357. 10.1186/s12864-019-5647-8.

[18] C. Alvarez-Carreño, P. Penev, A. Petrov, L. Williams, Fold Evolution before LUCA: Common Ancestry of SH3 Domains and OB Domains, Molecular Biology and Evolution 38 (2021). 10.1093/molbev/msab240.

[19] A.I. Azuaga, S.C. Atienza, Versatility of SH3 Domains in the Cellular Machinery, in: N. Kurochkina (Ed.), SH Domains: Structure, Mechanisms and Applications, Springer International Publishing, Cham, 2015: pp. 35–69. 10.1007/978-3-319-20098-9_3.

[20] J.Z. Lu, T. Fujiwara, H. Komatsuzawa, M. Sugai, J. Sakon, Cell wall-targeting domain of glycylglycine endopeptidase distinguishes among peptidoglycan cross-bridges, J Biol Chem 281 (2006) 549–58. 10.1074/jbc.M509691200.

[21] Y. Shen, I. Kalograiaki, A. Prunotto, M. Dunne, S. Boulos, N.M. I. Taylor, E. T. Sumrall, M. R. Eugster, R. Martin, A. Julian-Rodero, B. Gerber, P. G. Leiman, M. Menéndez, M. Dal Peraro, F. Javier Cañada, M. J. Loessner, Structural basis for recognition of bacterial cell wall teichoic acid by pseudo-symmetric SH3b-like repeats of a viral peptidoglycan hydrolase, Chemical Science 12 (2021) 576–589. 10.1039/D0SC04394J.

[22] U. Bodenhofer, E. Bonatesta, C. Horejš-Kainrath, S. Hochreiter, msa: an R package for multiple sequence alignment, Bioinformatics 31 (2015) 3997–3999. 10.1093/bioinformatics/btv494.

[23] D. Charif, J.R. Lobry, SeqinR 1.0-2: A Contributed Package to the R Project for Statistical Computing Devoted to Biological Sequences Retrieval and Analysis, in: U. Bastolla, M. Porto, H.E. Roman, M. Vendruscolo (Eds.), Structural Approaches to Sequence Evolution: Molecules, Networks, Populations, Springer, Berlin, Heidelberg, 2007: pp. 207–232. 10.1007/978-3-540-35306-5_10.

[24] K.P. Schliep, phangorn: phylogenetic analysis in R, Bioinformatics 27 (2011) 592–593. 10.1093/bioinformatics/btq706.

[25] S. Xu, L. Li, X. Luo, M. Chen, W. Tang, L. Zhan, Z. Dai, T.T. Lam, Y. Guan, G. Yu, Ggtree: A serialized data object for visualization of a phylogenetic tree and annotation data, iMeta 1 (2022) e56. 10.1002/imt2.56.

[26] M. Mirdita, K. Schütze, Y. Moriwaki, L. Heo, S. Ovchinnikov, M. Steinegger, ColabFold: making protein folding accessible to all, Nat Methods 19 (2022) 679– 682. 10.1038/s41592-022-01488-1.

[27] N. Guex, M.C. Peitsch, T. Schwede, Automated comparative protein structure modeling with SWISS-MODEL and Swiss-PdbViewer: a historical perspective, Electrophoresis 30 Suppl 1 (2009) S162–173. 10.1002/elps.200900140.

[28] L. Holm, DALI and the persistence of protein shape, Protein Sci 29 (2020) 128–140. 10.1002/pro.3749.

[29] P. García, C. Madera, B. Martínez, A. Rodríguez, J. Evaristo Suárez, Prevalence of bacteriophages infecting Staphylococcus aureus in dairy samples and their potential as biocontrol agents, Journal of Dairy Science 92 (2009) 3019–3026. 10.3168/jds.2008-1744.

[30] J. Valle, A. Toledo-Arana, C. Berasain, J.-M. Ghigo, B. Amorena, J.R. Penadés, I. Lasa, SarA and not σB is essential for biofilm development by Staphylococcus aureus, Molecular Microbiology 48 (2003) 1075–1087. 10.1046/j.1365-2958.2003.03493.x.

[31] S. Delgado, R. Arroyo, E. Jiménez, M.L. Marín, R. del Campo, L. Fernández, J.M. Rodríguez, Staphylococcus epidermidis strains isolated from breast milk of women suffering infectious mastitis: potential virulence traits and resistance to antibiotics, BMC Microbiol 9 (2009) 82. 10.1186/1471-2180-9-82.

[32] V. Martín, A. Maldonado-Barragán, L. Moles, M. Rodriguez-Baños, R.D. Campo, L. Fernández, J.M. Rodríguez, E. Jiménez, Sharing of bacterial strains between breast milk and infant feces, J Hum Lact 28 (2012) 36–44. 10.1177/0890334411424729.

[33] Harrell Jr, F, Hmisc: Harrell Miscellaneous, (2023). https://CRAN.R-pro-ject.org/package=Hmisc.

[34] Taiyun Wei, Viliam Simko, R package “corrplot”: Visualization of a Correlation Matrix, (2021). https://github.com/taiyun/corrplot.

[35] H. Wickham, ggplot2 elegant graphics for data analysis, 2nd ed., Springer International Publishing, 2016.

[36] B. Son, M. Kong, S. Ryu, The auxiliary role of the amidase domain in cell wall binding and exolytic activity of staphylococcal phage endolysins, Viruses 10 (2018) 284. 10.3390/v10060284.

[37] M. Landthaler, D.A. Shub, Unexpected abundance of self-splicing introns in the genome of bacteriophage Twort: Introns in multiple genes, a single gene with three introns, and exon skipping by group I ribozymes, Proceedings of the National Academy of Sciences 96 (1999) 7005–7010. 10.1073/pnas.96.12.7005.

[38] S. O’Flaherty, A. Coffey, R. Edwards, W. Meaney, G.F. Fitzgerald, R.P. Ross, Genome of Staphylococcal Phage K: a New Lineage of Myoviridae Infecting Gram-Positive Bacteria with a Low G+C Content, Journal of Bacteriology 186 (2004) 2862–2871. 10.1128/jb.186.9.2862-2871.2004.

[39] K. Vandersteegen, A.M. Kropinski, J.H.E. Nash, J.-P. Noben, K. Hermans, R. Lavigne, Romulus and Remus, Two Phage Isolates Representing a Distinct Clade within the Twortlikevirus Genus, Display Suitable Properties for Phage Therapy Applications, J Virol 87 (2013) 3237–3247. 10.1128/JVI.02763-12.

[40] D.R. Edgell, M. Belfort, D.A. Shub, Barriers to Intron Promiscuity in Bacteria, J Bacteriol 182 (2000) 5281–5289.

[41] D.H. Hall, Y. Liu, D.A. Shub, Exon shuffling by recombination between self-splicing introns of bacteriophage T4, Nature 340 (1989) 574–576. 10.1038/340574a0.

[42] D. Pinto, R. Gonçalo, M. Louro, M.S. Silva, G. Hernandez, T.N. Cordeiro, C. Cordeiro, C. São-José, On the Occurrence and Multimerization of Two-Polypeptide Phage Endolysins Encoded in Single Genes, Microbiology Spectrum 10 (2022) e01037–22. 10.1128/spectrum.01037-22.

[43] I. Abaev, J. Foster-Frey, O. Korobova, N. Shishkova, N. Kiseleva, P. Kopylov, S. Pryamchuk, M. Schmelcher, S.C. Becker, D.M. Donovan, Staphylococcal phage 2638A endolysin is lytic for Staphylococcus aureus and harbors an inter-lytic-domain secondary translational start site, Appl Microbiol Biotechnol 97 (2013) 3449– 3456. 10.1007/s00253-012-4252-4.

[44] U. Dionne, É. Bourgault, A.K. Dubé, D. Bradley, F.J.M. Chartier, R. Dandage, S. Dibyachintan, P.C. Després, G.D. Gish, N.T.H. Pham, M. Létourneau, J.-P. Lambert, N. Doucet, N. Bisson, C.R. Landry, Protein context shapes the specificity of SH3 domain-mediated interactions in vivo, Nat Commun 12 (2021) 1597. 10.1038/s41467-021-21873-2.

[45] A. Grundling, O. Schneewind, Cross-linked peptidoglycan mediates lysostaphin binding to the cell wall envelope of *Staphylococcus aureus*, J Bacteriol 188 (2006) 2463–72. 10.1128/JB.188.7.2463-2472.2006.

[46] L.S. Gonzalez-Delgado, H. Walters-Morgan, B. Salamaga, A.J. Robertson, A.M. Hounslow, E. Jagielska, I. Sabała, M.P. Williamson, A.L. Lovering, S. Mesnage, Two-site recognition of Staphylococcus aureus peptidoglycan by lysostaphin SH3b, Nat Chem Biol 16 (2020) 24–30. 10.1038/s41589-019-0393-4.

[47] R. Diez-Martinez, H.D. De Paz, E. Garcia-Fernandez, N. Bustamante, C.W. Euler, V.A. Fischetti, M. Menendez, P. Garcia, A novel chimeric phage lysin with high *in vitro* and *in vivo* bactericidal activity against *Streptococcus pneumoniae*, J Antimicrob Chemother 70 (2015) 1763–73. 10.1093/jac/dkv038.

[48] M.J. Loessner, K. Kramer, F. Ebel, S. Scherer, C-terminal domains of *Listeria monocytogenes* bacteriophage murein hydrolases determine specific recognition and high-affinity binding to bacterial cell wall carbohydrates, Mol Microbiol 44 (2002) 335–49. 10.1046/j.1365-2958.2002.02889.x.

[49] A. Gouveia, D. Pinto, J.M.B. Vítor, C. São-José, Cellular and Enzymatic Determinants Impacting the Exolytic Action of an Anti-Staphylococcal Enzybiotic, International Journal of Molecular Sciences 25 (2024) 523. 10.3390/ijms25010523.

