## Supplementary Figure for "Diversity, structure-function relationships and evolution of cell wall-binding domains of staphylococcal phage endolysins"

**Supplementary Table S1**. Pairwise similarity distance matrix of the 816 endolysin sequences retrieved from PhaLP (provided as a separate file).

**Supplementary Table S2**. Curated set of 182 staphylococcal endolysins (provided as a separate file).

**Supplementary Table S3**. Pairwise similarity matrix based on a DALI 3D structural comparison of the 3D predictions obtained for a selected subset of 42 representative CBD repeats (provided as a separate file).

**Supplementary Table S4**. Summarized results corresponding to the specificity profiles of the 24 selected CBDs versus the collection of 24 staphylococcal strains.

**Supplementary File S1**. 3D structures of the trimmed CBDs used for the DALI structural alignment detailed in Supplementary Table S3 and Figure 2 (provided as a separate file).

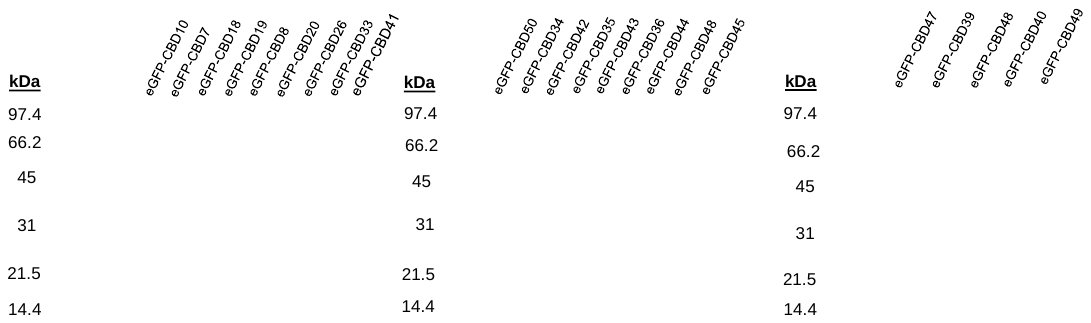

**Supplementary Figure S1.** SDS-PAGE of the purified eGFP-CBD fusions. The expected molecular weight of each protein (in kDa) is shown on top of each band.

**Legend:** Endolysin in dataset; incomplete endolysin fragment; insertion/intron; HNH endonuclease ; staphylokinase

>A0A0U1WF03_alternativeID:AHB79986_LysK_phage-K

ATGGCTAAGACTCAAGCAGAAATAAATAAACGTTTAGATGCTTATGCAAAAGGAACAGTAGATAGCCCTTACAGAGTTAAAAAAGCTACAAGTTATGACCCATCATTTGGTGTAATGGAAGCAGGAGCCATTGATGCAGATGGTTACTATCACGCTCAGTGTCAAGACCTTATTACAGACTATGTTTTATGGTTAACAGATAATAAAGTTAGAACTTGGGGTAATGCTAAAGACCAAATTAAACAGAGTTATGGTACTGGATTTAAAATACATGAAAATAAACCTTCTACTGTACCTAAAAAAGGTTGGATTGCGGTATTTACATCCGGTAGTTATGAACAGTGGGGTCACATAGGTATTGTATATGATGGAGGTAATACTTCTACATTTACTATTTTAGAGCAAAACTGGAATGGTTATGCTAATAAAAAACCTACAAAACGTGTAGATAATTATTACGGATTAACTCACTTCATTGAAATACCTGTAAAAGCAGGAACTACTGTTAAAAAAGAAACAGCTAAGAAAAGCGCAAGTAAAACGCCTGCACCTAAAAAGAAAGCAACACTAAAAGTTTCTAAGAATCACATTAACTATACAATGGATAAACGTGGTAAAAAACCTGAAGGAATGGTAATACACAACGATGCAGGTCGTTCTTCAGGACAACAATACGAGAATTCATTAGCTAATGCAGGTTATGCTAGATACGCTAATGGTATTGCTCATTACTACGGCTCTGAAGGTTATGTATGGGAAGCAATAGATGCTAAGAATCAAATTGCTTGGCACACGGGTAAATAAATTGCCCTGTTATACAGTAATGTATAATGAAAGTTCTTTTAATTGACTGGGAGGCTAAGGCTATCAAGCTATGCTAATCAGCATCCAAGAATATTGTTTTAAAAAACTACTTGACAACATAATAACTTTCCTATATACTTAAGTAAAGGAGATGTTATTATGGAAAAGAAATTAAATGAAATACCTGGATTAGAAATATATGAAAATTACACTATTACTGATAAAGGAGAAGTAATATCTTATAAAGGTAAAGAGCCTAAAAAGTTAAAACTTCAAAAAAATAACAAGGGTTACTTGTTTGTAAGGTTACGATACCATTCACCTAAAATACATCGTTTAGTTGCTATGGCTTTTATACCTAATCCTGATAATAAAGAACAAGTTAACCATTTAAATGGTAAAAATGATAATAGTGTAGGAAATTTAGAATGGGTTTCTAATTCCGAGAACAGAGAACATGCAATAAAGACAGGATTAAAAAATGAAATAAATTATAATATAGCTCAGTATGACTTAGAAGGTAATTTATTGAATGTCTTTTACACAGCTCAAGAGGCTTTAGAGTTCTTAGGTATTTCTAATAAAAGAAGTGGTAATATAGGAAGATGTATCAAAGGAGAGAGAAAAACAGCCTACGGATACATTTGGAAACAATATTAAGGTTCAACGACTATCCCTCGGACGAGTGCCAATAAAAATAAATCGTCAATAGGAGTAGGGCTCAAGTTAATGGAGTGGGTGAGAATCCCTTAAATCGAAATGGAGAACCCCTAACAAAATTAGGGTGAAGATATAGTCTAGTCTTATAGGAAACTATAAGGAGTTCATAAGAGAACCGGTAAAGGAGTTACGTCTTTTATTGAATAATCGGATGGAACAGGAGCAAACTCAGGTAACTTTAGATTTGCAGGTATTGAAGTCTGTCAATCAATGAGTGCTAGTGATGCTCAATTCCTTAAAAATGAACAAGCAGTATTCCAATTTACAGCAGAGAAATTTAAAGAATGGGGTCTTACTCCTAACCGTAAAACTGTAAGATTGCATATGGAATTTGTACCAACTGCCTGTCCTCACCGTTCTATGGTTCTTCATACAGGATTTAATCCAGTAACACAAGGAAGACCATCACAAGCAATAATGAATAAATTAAAAGATTATTTCATTAAACAAATTAAAAACTACATGGATAAAGGAACTTCAAGTTCTACAGTAGTTAAAGATGGTAAAACAAGTAGCGCAAGTACACCGGCAACTAGACCAGTTACAGGTTCTTGGAAAAAGAACCAGTACGGAACTTGGTATAAACCGGAAAATGCAACATTTGTCAATGGTAACCAACCTATAGTAACTAGAATAGGTTCTCCATTCTTAAATGCTCCAGTAGGCGGTAACTTACCGGCAGGGGCTACAATTGTATATGACGAAGTTTGTATCCAAGCAGGTCACATTTGGATAGGTTATAATGCTTACAACGGTAACAGAGTATATTGCCCTGTTAGAACTTGTCAAGGTGTTCCACCTAATCAAATACCTGGCGTTGCCTGGGGAGTATTCAAATAG

>H9A141

ATGAAAACACAATCTCAAATCAATAAACGTTTAAGAGATTATAAAAACGGTGTAGTAGATAGTCCATACAGAGTTAAACATTGGACGAGTTATGACGCTTCCTTTGGTGCTATGGAACCAGGTTGCATTGATAAAGACCGTGCTTATCACGCACAGTGTATGGATTTGGCGATAGATTATGTAATGTGGTTAACTGATAATCAAACAGAGATGTGGGGCGATGCTAAAAGCTCTATAAAAAACAAATTCCCTAAAGGGTGGAAGATTGTAGAAAACAAACCGTCAACGATACCCAAAAAAGGTTGGATAGCTGTATATACAGCTGGAACCTATTCACGTTATGGGCACATTGGTATTGTATATAATGGTGGTAATACGAATAGCTTCCAAATTTTAGAACAAAATTGGAATGGCTGGGCTAATAAAAAACCTAGCTTACGATGGGATAACTATTATGGTTTAACACATTTTATAGTTCCACCGGTAGCGAAAGAAATAGAAGAACCTAAGAAAGATGTAAAATCAGCTCCTAAACAGTCAGTTAAGAAAAATAGTAGTATCAAAGTTAACACGCATCATATAAAAGGTTGGACTATGACTAAAAGAGGTCGTAAACCTAAAGGTGTATAATAGTGCAAATCCCCTTCATATATAGTATAATACCTGTATAATGACTTAGGAGAGGTATTATATGAAAGAGATATGGAAAGATATAAAAGGTTATGAAAATTATTATATGATTTCTAATACAGGTAAGGTTAAATCTTTAAAGAGAGTTATTAAGCACGGCGGTTCAAAGACTAAAACGCTCCCCGAAACTATTTTAAAACCAAATAAAGTTAATTTCGGGTATTTACAAGTTACTTTAAACAAAAACGGAAAAAGAAAGTGTAAGTATATTCACAATTTAGTTATGGAAACTTTTGTTGGTCCAAAAAAAATAGGATACGAAGTGAACCATAAGAACGAAAACAAAGAGGATAATTCTTTGGAAAATTTAGAATATTTGACTTGTAAAGAAAATAACAATTACGGTACTAGGATACAAAGATATGTTGAAAAGGCTAAAAATGGAAAACGTAGCAAAAAAATTAAAGGAACTCATATAATAACTGGAGAAATCATATACTTCCCTTCTATATCTGAAGCTAGAAGACAAGGATATGGAAAACATATATCTGAAGTTTGCAGAGGTTATAGAAATCACTGCAATAATTACAAATGGGAATTTGTATAAGCACTGCATATATAGCGATGTATATGTACTAAAAACATAAATTGCTTTGAAGAACCTTAGAGCCTTAATACCACAATAAAGCTGAAAAGACTTTTTGACGGTTTAACAATTTTAAGGATTGGTTTGTTTAGCAGCACTACCCCTAAGTGGAGACATATGGGGAATGTTCAACGACTAGAGGATTGCGTCCTCGTAGGTTTCAAGTGAAATCGAAAGAAGAACTATCTCACGTAGATAGAGACATATAGTCTGGTCACGTCTTGTAATGAGAGTGCTAGGAATTAACCTAGAGTTAGACTAACGACCTAACAAAACAAAACGTAGTTATCCATAACGATGCCGGTACAATGAATTCTAAACAATACTATAACAATTTGGTAAACGCTGATTACAATAGGTTAGCACGAGGTATAGCTCATGCATACGCTGATAGAAACGGTATTTGGGAAGCTATATCAGAAGATAGAATTGCTTGGCATGTTTCTGATGGCGTTCAACCAGGTTCAGGTAATTTTGAAACTTATGGAATTGAAGTTAATCAATCAATGTATGTAAGTGATAAAGATTTCCTTAAAAATGAACAAGCAGCTCTTAAATTCGCAGCGCATAAACTTAAAAAGTGGGGGTTACCAGCTAACAGAAATACTGTTCGTTTACACAACGAATTTAGTTATACAGCTTGTCCTCATCGTTCAGCTAAATTGCATACTGGTATTGATCCAACAAAACAAGCATGGACTAAAGCGACACAACTTAAGTTAAAAGATTACTTCATTAAACAAATTAGGGCGTATATGAAAGGTGATACACCTAAGGTTACTACGGTCAAAAATAAACCTGGTAGTGCTTCTACTCCTGCTAACAGACGAGATATGAACGGTTGGAAAATCAATAAGTATGGAACTTATTACAAATCAGAAGTAGCTCATTTTACGCCAAACACGCCTATTAAAACTCATTATGTTGGACCGTTTAGAAGTTGTCCTGTGAGCGGTGTATTACAACCCGGACAAACGGTAAGATACGATACCGTATGTAAACAAGATGATCACGTTTGGATTAGTTACACAGCCTACAATGGCAAAGATGTGTGGTTAGCAGTAAGAACATGGGACAAAAATACAGATAGTTTAGGTAAGTTGTGGGGGACAATTAAATAA

>A0A345AQJ3

ATGAAAACATACAGTGAAGCAAGAGCAAGGTTACGTTGGTATCAAGGTAGATATATTGATTTTGACGGTTGGTATGGTTACCAATGTGCAGATTTAGCAGTTGATTACATTTATTGGTTGTTAGAAATTAGAATGTGGGGAAATGCAAAAGATGCAATCAATAACGATTTTAAAAACATGGCAACAGTATATGAAAACACACCATCGTTTGTTCCACAAATAGGTGATGTGGCTGTATTTACCAAAGGAATATATAAACAATACGGTCATATTGGTTTAGTGTTTAATGGTGGTAATACAAACCAATTTTTAATTTTGGAACAGAACTATGACGGTAACGCAAATACGCCTGCAAAGTTACGTTGGGATAATTATTACGGCTGTACTCACTTTATTAGACCTAAGTATAAAAGTGAGGGCTTAATGAATAAGATCACAAATAAAGTTAAACCACCTGCTCAAAAAGCAGTCGGTAAATCTGCAAGTAAAATAACAGTTGGAAGTAAAGCGCCTTATAACCTTAAATGGTCAAAAGGTGCTTATTTTAATGCGAAAATCGACGGCTTAGGTGCTACTTCAGCCACTAGATACGGTGATAATCGTACTAACTATAGATTCGATGTTGGACAGGCTGTATACGCGCCTGGAACATTAATATATGTGTTTGAAATTATAGATGGTTGGTGTCGCATTTATTGGAACAATCATAATGAGTGGATATGGCATGAGAGATTGATTGTGAAAGAAGTGTTTTAATTCTTAGGTTAAAATGTTAAATATTTGTTAATTATTTTTTAATGTAAGTTTAGTTTCTTTTAATATTTTATTGATTTTTAATATTTTCTCAATATAAAATGAAGTTGTTGATATTTATCATCTTAAATAAGGGTGTTAGCTATAAAAAGAGATAAATAAAAACAAATATATTATATTTGGAGGAAGCGCCATGCTCAAAAGAAGTTTATTATTTTTAACTGTTTTATTGTTATTATTCTCATTTTCTTCAATTACTAATGAGGTAAGTGCATCAAGTTCATTCGACAAAGGAAAATATAAAAAAGGCGATGACGCGAGTTATTTTGAACCAACAGGCCCGTATTTGATGGTAAATGTGACTGGAGTTGATGGTAAAGGAAATGAATTGCTATCCCCTCGTTATGTCGAGTTTCCTATTAAACCTGGGACTACACTTACAAAAGAAAAAATTGAATACTATGTCGAATGGGCATTAGATGCGACAGCATATAAAGAGTTTAGAGTAGTTGAATTAGATCCAAGCGCAAAGATCGAAGTCACTTATTATGATAAGAATAAGAAAAAAGAAGAAACGAAGTCTTTCCCTATAACAGAAAAAGGTTTTGTTGTCCCAGATTTATCAGAGCATATTAAAAACCCTGGATTCAACTTAATTACAAAGGTTGTTATAGAAAAGAAATAAAACAAAATAGTTGTTTATTATAGAAAGCAATGTCTTGCTTGAATATGTGTAGTGAAAATTATCTTTCATCAAATTCTCATTCATGCACGAATGGCTCTTCCCCACCTAATCAGATATTAGGTGACTTATGGGGAGAAATCAGTTAGGATGAAAAAGTGGATAATCCTTTTTTTAGGCAGGTACTTCGGTACTTGCCTATTTTTTTATGTTATAATCTTTCTAGACGTATTCAAAGGACGTCTTTTTAGAACGTATGTTATAGCTAGCTTTCGGGCTAGTTTTTTGCTATGATGCGTTACACATGCATCAACTATTTACATCTATCTTTGTTCACCTAAGCATGTCACTGGGTGTTTTTTTCTTACGATAGAGAGCATAGTTTTCATACTACTCCCCGTAGTATATATGACTTTAGCATTCCCGTATAACAGTTTACGGGGTGCTTTTTATGTTATACTTAACTGTATATAGTAGGAGTGAACTATATAGCCTGTTAAGTGGCCTAGTAACCTAACACTTATCCTGCAATTGATATCCTTTTTGCCCTTCACTCGATACATATATCTCAACAACATAGAAATATTACAGTCGCTACACCGCATCTTAAATGGTGTGGTTATTTTTATTGGAAGTGTGTATCAGGTATCAGTAATGTTAAAACACCAGCTAAAAATGAAAAGAATTCACCAGTGCCAGCAGGTTATACACTTGATAAGAATAATGTGCCTTATAAAAAAGAGACTGGTAATTACACAGTTGCCAATGTTAAAGGTAATAACGTAAGGGACGGCTATTCAACTAATTCAAGAATTACAGGTGTATTACCTAATAACGCAACAATCAAATATGACGGCGCATATTGCATCAATGGGTATAGATGGATTACTTATATTGCTAATAGTGGACAACGTCGCTATATTGCGACAGGAGAGGTAGATAAAGCAGGTAATAGGATAAGTAGTTTTGGTAAGTTTAGCGCAGTTTGA

>Q4ZC44

GTGACAAGAACTTACCCGACTGTGATTTTAAAACCAAATAAAGTAGCATTTGATTATTTTCAAGTCACTTTAAATAAAAATAGCATAAGAAAATCAAGATACGTTCACAATTTGGTTATGGAAGCATTTGTCGGAGAGAAGAAGTTAGGTTATGAAGTTAATCATAAAAACGAAGATAAAAGTGATAATCGTCTAGAAAACCTAGAATATATAACTTCTAAAGAAAATAATAATTATGGAACTAGAATCGAAAGATCTATTAAAAAATCTACAAATGGTAAAAGAAGTAAGAAAATCAAAGGAACTCATATAATTACTGGAGAAGAAGTTTATTTTCCATCTATTGCTGAAGCAAAAAGACAGGGATATGGTAATCATATTAGCGACGCAATACGTGGTAAAAGAAATCACTGTTATAAATATAAATGGGAATTCATTTAAGCGCTGCATATATAGCGATGTATATGTACTAAAAACATAAATTGCTTTGAAGAACCTTAGAGCCTTAATACCACAATAAAGCTGAAAAGACTTTGTGAAGGTTTAACAAGTTTAAGGATTGGTTTGTTTAGCAGCACTACTCCTAAATACTTCTTAGTACAAGGAGAATGTTCAACGACTAGAGGATTGCGTCCTCGTAGGGTTCAAGTGAACTCGAAAGAAGAACTATCTCATGTAGATAGAAACATATAGTCTGGTCACGTCTTGTAATGAGAGTGCTAGGGATAAACCTAGAGTTAGATTAACGACCTAACAAAACAAACGTAGTTATACACAACGACGCAGGAAGCAAAGGAGCGACTGCTGAAGCATATCGTAATGGGTTAGTTAATGCGCCTTTATCGAGACTAGAGGCAGGTATTGCGCATAGTTATGTATCAGGTAACACAGTGTGGCAAGCCTTAGATGAATCACAAGTAGGTTGGCATACTGCTAACCAATTAGGCAATAAATATTATTACGGTATTGAAGTGTGTCAATCAATGGGAGCGGATAATGCGACGTTTTTAAAAAATGAACAGGCGACTTTCCAAGAATGCGCTAGATTGTTGAAAAAATGGGGATTACCAGCAAACAGAAATACAATCAGATTACACAACGAATTCACTTCAACATCATGCCCACACAGAAGCTCAGTATTGCACACTGGTTTTGACCCAGTAACTCGTGGCCTATTGCCGGAAGATAAACAATTACAACTTAAAGACTACTTTATCAAGCAAATTAGAGTGTATATGGACGGTAAGATACCAGTTGCCACTGTCTCTAATGAGTCAAGCGCTTCAAGTAATACAGTTAAACCAGTTGCGAGTGCATGGAAACGTAATAAATATGGTACTTACTACATGGAAGAAAGTGCTAGATTCACAAACGGTAATCAACCAATCACTGTAAGAAAAATAGGACCATTCTTATCATGCCCGGTAGCTTACCAATTCCAACCTGGTGGATATTGTGATTATACAGAAGTGATGTTACAAGATGGTCATGTTTGGGTAGGATATACATGGGAGGGGCAACGTTATTACTTGCCTATTAGAACATGGAATGGTTCTGCCCCACCTAATCAGATATTAGGTGACTTATGGGGAGAAATCAGTTAG

>A0A6F8Z5Y5

ATGTTGATAACAAAAAACCAAGCAGAAAAATGGTTTGATAATTCATTAGGGAAGCAGTTCAATCCTGATTTGTTTTATGGATTTCAGTGTTACGATTACGCAAATATGTTTTTTATGATAGCAACAGGCGAAAGGTTACAAGGTTTATACGCTTATAATATTCCATTTGATAATAAAGCAAGGATTGAAAAATACGGGCAAATAATTAAAAACTATGATAGCTTTTTACCGCAAAAGTTGGACATTGTCGTTTTCCCGTCAAAGTATGGTGGCGGAGCTGGACATGTTGAAATTGTTGAGAGCGCAAATCTAAACACTTTCACATCATATGGCCAAAATTGGAATGGTAAAGGTTGGACAAATGGCGTTGCGCAACCTGGTTGGGGTCCCGAAAGCTGTTACAAGACATGTTCATTATTACGATGACCCAATGTATTTTATTAGATTAAATTTCCCAGATAAAGTAAGTGTTGGAGATAAAGCTAAAAACGTTATTAAGCAAGCAACTGCCAAAAAGCAAGCAGTAATTAAACCTAAAAAAATTATGCTTGTAGCCGGTCATGGTTATAACGATCCTGGAGCAGTAGGAAACGGAACAAACGAACGCGATTTTATCCGTAAATATATAACACCAAATATCGCTAAGTATTTAAGACATGCAGGTCACGAAGTTGCATTATATGGTGGCTCAAGTCAATCACAAGACATGTATCAAGATACTGCATACGGTGTTAATGTAGGAAATAATAAAGATTATGGATTATATTGGGTTAAATCACAGGGGTATGACATTGTTCTAGAGATTCATTTAGACGCAGCAGGAGAAAATGCAAGTGGTGGGCATGTTATTATCTCAAGTCAATTCAATGCAGATACTATTGATAAAAGTATACAAGATGTTATTAAAAATAACTTAGGACAAATAAGAGGTGTAACACCTCGTAATGATTTACTGAACGTTAATGTATCAGCAGAAATAAATATCAATTATCGTTTATCTGAATTAGGTTTTATTACTAATAAAAAAGATATGGATTGGATTAAGAAGAATTATGACTTGTATTCTAAATTAATAGCTGGTGCAATTCATGGTAAGCCTATAGGTGGTTTGGTAGCTGGTAATGTTAAAACATCAGCTAAAAACCAAAAAAATCCACCAGTGCCAGCAGGTTATACACCCGATAAAAATAATGTACCGTATAAAAAAGAAACTGGTAATTACACAGTTGCCAATGTTAAAGGTAATAACGTAAGGGACGGCTATTCAACTAATTCAAGAATTACAGGTGTATTACCTAATAACGCAACAATCAAATATGACGGAGCATATTGCATCAATGGGTATAGATGGATTACTTATATTGCTAATAGTGGACAACGTCGTTATATTGCTACAGGAGAGGTAGACAAGGCAGGTAATAGAATAAGCAGTTTTGGTAAGTTTAGTGCAGTTTGA

>A0A060AB53

GTGGCAACTTTAACGCATAAAGAAGCCGTAGAATATGTTAAATCATTAGAAGGAAAATATGTTGACTTTGATGGTTGGTATGGGTTAATTGGTAGCCCATGTAAAACTTTTTGAATTGCTGGAAACCCCTAACGTAAAGACGAGGGCAATCAGCAGCGAAGCTCACATGGTAACAGTGTGTGAACGTTCAACGACTATGTACTATCAATTGATAGGCAGCGGAAAGCGTTTGACAAATATATACTCCTGTGATATAATGTTTTAATATGAAAATTTTGTCAAATAATGATATAGTCTGGTCTCATATGAAAATATGAGGCTTTCTTTATGAAAGCTATTAAATGTTATACAATATTATATAGAAATATATAATAGGAAAGGCATTTAATAAAACAAAAAACGACCAATGTTTTGATTTAGCTAACAAATATTGGAATAAATTATTTGGTGGACAATTAAAAGGTCAAGGCGCAGCTGATATTCCTAATGTTAATGATTTTAGTGGAAAAGCTACTGTTTATCAAAATACAACAAGTTTCTTAGCAAAACCAGGCGACATTGTTGTATGGGGTAGAAACTTTGGTCAAGGATATGGTCATGTAGCAGTTGTTGTTGAAGCAACATTGGATTACATTGTTGTAATTGAGAATAACTGGTTAGGTGGTGGATGGACTAGTGGACCGGCACAAGGTGGTACTGGTTGGGAAAAAGCAACTAGACGCCGTCACAATTACGAGTTTTCTATGTGGTTCATCAGACCAAAATACAAAGACGAGAAAAAAATTACTAATTATTCTGCTAAAACTAAAACGAAAAAACCAGCTGCTAAAAAGAAAAAAGCTAAAGAAATAAAACACATTAAAGATACAGTTAACGGTTATAAATTACCAAAACGTAACGGTAAATTAAAAGGTGTCGTGTAAAATGCCTATAAATTAATTGACTTTTTTATTTCTTTATGATATAATGACTATATATTTATATATATAGGAGGATATTATATGAAAGAAATTTGGAAAGATGTTCCTAATTATGAAGGAAAATATCAAGTGTCTAATAAAGGAAGAGTTAAATCAATTAAAAGAACAGTAAAACATAACGGGAGTAAAACTAGGACATTTCCAGAGAAGATTTTAAAACCAAACAAAGTGTCTTTTGATTATTTACAGGTCACATTATATAATAATGGTAAAAGAAAATGTAGATATATACACAATTTAGTTATGGAATCTTTTATTGGTAAGAAGCCAAACGGTTATGAAGTTAACCATATAGACGAAGATAAATCTAATAATCAATTAGAAAATTTAGAATATATAACAAGAAAAGAAAACAATAATTATGGAACTCGAATAGAACGGATGATTAAATCAAACGTTAATGGTAAAAAGAGTAAAAAAGTCAAAGGGACCCATTTAAAAACAGGAGAAATAGTTATTTTTCCATCAATATCCGAAGCGCAAAGACAAGGATATGGACCAAAAATATCAGAAGTTTGCAATGGAAATAGAAATCATTCCGGTGGTTATAAATGGGAATTTATAAAATAGCACTGCATAGAAAGGAAACTACTATGTACTAAAAACATAAATTGCTTAGAAGAACCTTAGAGCCTTAATGCCACAATATAGCTGAAAAGACTGTATGAAGGATTAAAAAGTTTAAGGATTGGTTTGTTTAGCATCGCTACTCCTAAGTCTTTTAGATATGGAGAACGTTCAACGACTAGACGATTGGCTCGTCGTAGGGTTTAAGTAAACTCGAAAGAAGAACTACCTAAGTATTTTATATATGGTAGAAAGATATAGTCTGGACACGCCTTTAATAGGGGTGCTAGGAATCGACCTAGAATTAGATTAACGACCTAATGAAACGTTTCGATACACAATGATGCCGGAAGTAAATATGCTACAGCAGAAGCTTATCGTAATGGATTAGTTAATGCTCCATTATCAAGACTTGAAGCCGGTATTGCGCATGCTTACGGCTCCGGTAATACAATTTGGCAAGCATTAGATAAATCACAAGTAGGCTGGCACACAGCAAATCCTGTTGGTAACAACGGTTATTATGGTTATGAAGTTTGTCAATCTATGGGTGCAGATGACAAAACATTCTTAGCAAATGAACAACTAGTATTCCAAGATGCTGCTAGACTTTTAAAAGAAGAAGGACTACCAGCAAACAGAAATACTGTTCGTCTACATTGTGAATTTGTTCGTACTTCATGTCCACATAGAAGTGCTAAACTACACACAGGATTTGACCCTGTTACTCAAGGTTTATTACCTAAAGACAAACAGTTACAATTAAAAGATTATTTCATTAAACAAATCAGACAATATATGGATGGTAAAATACCTACTGCAACAGTTGTTAAAAGTACAAGTGCTTCTAGTAACACTGTTAAACCAGTAGCATCTGGTTGGAAACGTAATAATTACGGCACTTATTATAAATCAGAAAATGCAACTTATAGCAATGGTAACACACCAATTATCACACGTACAGTAGGACCATTTAGAAGCTGTCCTCAAGCAGGATTGTTACCAGCTGGAGCAACTATTGTATATGATACAGTTTGTTTACAAGACTATCATGTTTGGGTAAGTTATGTAACTAACAAAGGTTACAGAGTATGGCTACCTATTAGAACATGGAATGGTGTAGCTCCTGGAAACGCAGGATATGCTGTAGGACCATTATGGGGATATATCAGCTAA

**Supplementary Figure S2.** Genomic structure of the surrounding region of the endolysin for some selected examples indicated at Figure 3B, including an experimentally proven intron-containing endolysin gene (from staphylococcal phage K).

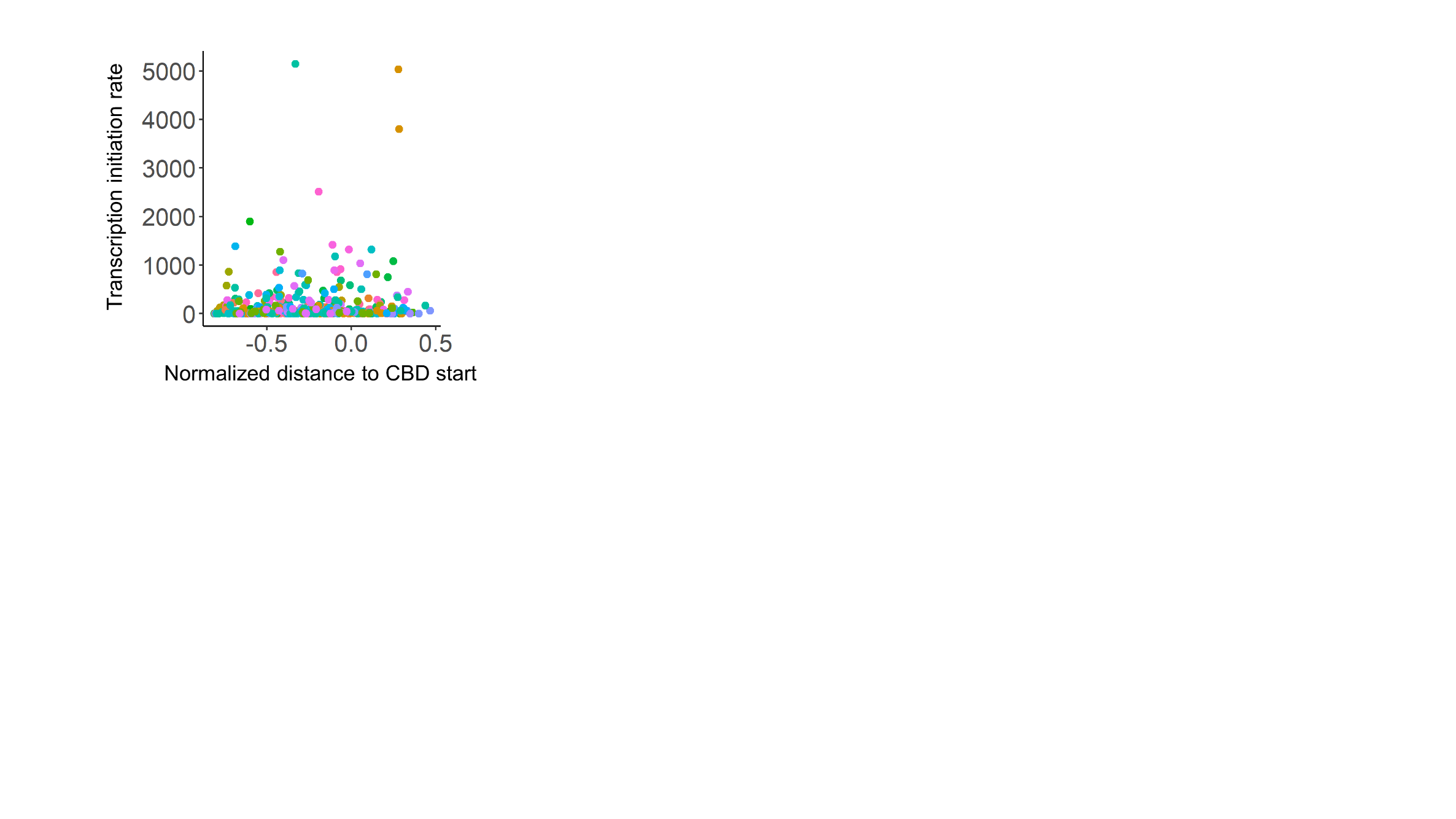

**Supplementary Figure S3.** In-frame RBS predictions as obtained from RBS Calculator in Predict Mode using the mRNA sequences of 42 staphylococcal endolysins. Dots of different colors are different endolysins. Virtually no significant predictions were found as almost no datapoints were clearly above (≥ one order of magnitude) the background values obtained for each sequence.

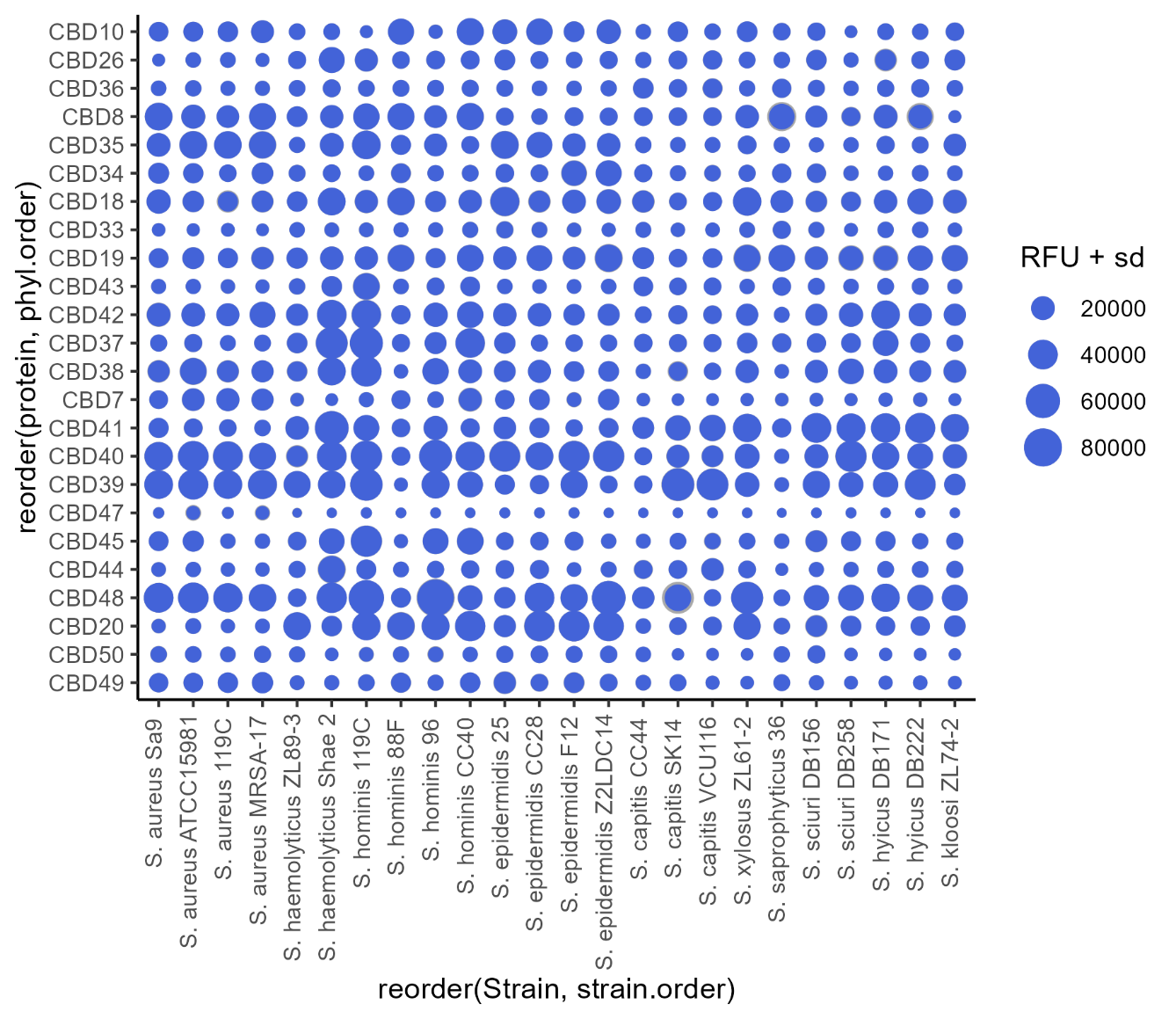

**Supplementary Figure S4.** Raw fluorescence measurements for each CBD/strain pair. The size of the blue spot represents the fluorescence intensity, and the surrounding grey shade is the standard deviation.
